# Tubulin mRNA stability is sensitive to change in microtubule dynamics caused by multiple physiological and toxic cues

**DOI:** 10.1101/533224

**Authors:** Ivana Gasic, Sarah A. Boswell, Timothy J. Mitchison

## Abstract

The localization, mass, and dynamics of microtubules are important in many processes. Cells may actively monitor the state of their microtubules and respond to perturbation, but how this occurs outside mitosis is poorly understood. We used gene expression analysis in quiescent cells to analyze responses to subtle and strong perturbation of microtubules. Genes encoding α-, β, and γ-tubulins, but not δ- or ε-tubulins, exhibited the strongest differential expression response to microtubule-stabilizing versus destabilizing drugs. Q-PCR of exon versus intron sequences confirmed that these changes were caused by regulation of tubulin mRNA stability and not transcription. Using tubulin mRNA stability as a signature to query the GEO database, we find that tubulin genes respond to toxins known to damage microtubules. Importantly, we find many other experimental perturbations, including multiple signaling and metabolic inputs that trigger tubulin differential expression, suggesting their novel role in the regulation of microtubule cytoskeleton. Mechanistic follow up confirms that one important physiological signal, phosphatidylinositol-4,5-bisphosphate 3-kinase (PI3K) activity, indeed regulates tubulin mRNA stability via changes in microtubule dynamics. We propose that that tubulin gene expression is regulated as part of many coordinated biological responses, with wide implications in physiology and toxicology. Furthermore, we present a new way to discover microtubule regulation using genomics.

## Introduction

α- and β-tubulin proteins form obligate heterodimers which, through nucleating activity of γ-tubulin, polymerize into microtubules. Microtubules physically organize eukaryotic cells, serve as platforms for intracellular transport and signaling, and power cell division^1^. The localization, mass, and dynamics of microtubules must be precisely tuned for microtubule cytoskeleton to execute its many functions. To ensure this, it seems likely that cells actively monitor the state of their microtubules and generate responses specific to perturbation. During mitosis, the Spindle Assembly Checkpoint (SAC) monitors the microtubules of the mitotic spindle and controls anaphase progression^2^. Efficiency of microtubule-targeting cancer chemotherapy has been associated with activation of the SAC and mitotic arrest^3^. However, evidence is growing that microtubule-damaging chemotherapy also triggers active signaling responses in interphase cells, which may contribute to cell death and tumor shrinkage^4^. Specifically, drug-induced microtubule damage in interphase has been associated with changes in cell cycle progression^5-7^, activation of mitogen-activated protein kinase (MAPK) signaling pathways^8-10^, and activation of guanine nucleotide exchange factors for Rho-family GTPases^11^. These data show that interphase cells respond to microtubule damage through some signaling pathways. However, a comprehensive view is missing, leaving a deliberate surveillance mechanism of the microtubule cytoskeleton in interphase cells speculative.

One route to discovering if, and how, interphase cells sense and respond to changes in microtubule cytoskeleton is to measure differential gene expression as a function of drug-induced microtubule damage. Naively, one might expect microtubule-stabilizing and destabilizing drugs to have opposite effects on expression of genes involved in monitoring microtubule damage. Consistent with this hypothesis, microtubule destabilization suppresses, while stabilization activates, synthesis of α- and β-tubulin^12-14^. This mechanism, known as tubulin autoregulation, is a negative feedback loop that involves indirect co-translational regulation of the stability of mature spliced tubulin mRNA by unpolymerized tubulin^12-14^. It remains unknown if tubulin autoregulation is part of a larger differential gene expression program. Several studies analyzed the effect of one or multiple microtubule-targeting drugs on gene expression using genome-wide methods^15-17^. One large-scale study found similar, rather than opposite, effects of microtubule-stabilizing versus destabilizing drugs^15^. A plausible explanation is that both stabilizers and destabilizers cause mitotic arrest, leading to common downstream effects on gene expression. The resulting changes in gene expression are thus likely to reflect cell cycle regulation more than microtubule-specific signaling.

We sought to discover specific gene networks that respond differentially to microtubule-stabilizing versus destabilizing drugs. To avoid a mitotic arrest signature, we used quiescent cells. Our study revealed several differential gene expression (DGE) signatures that clearly exhibited opposite responses to microtubule destabilization versus stabilization. We found that α, β and γ-tubulin genes exhibit the strongest differential regulation. Using tubulin DGE as a query in bioinformatics analyses, we discovered multiple physiological inputs that changed tubulin gene expression, including the PI3K signaling pathway. Follow up mechanistic analysis confirmed that PI3K activity regulated tubulin gene expression via changes in microtubule stability, leading to autoregulation of tubulin mRNAs. Our study reveals a new role of tubulin gene expression regulation as part of concerted responses of cells to multiple physiological, or damaging inputs.

## Results

### Gene expression profiles of microtubule perturbation in quiescent cells

To identify interphase-specific responses to microtubule damage, we first established conditions of healthy quiescence in a reference cell-type for cell biology studies. RPE1 hTert cells were grown to, and maintained as, confluent cultures for 5 days prior to treatment with microtubule-destabilizing and stabilizing drugs. Cell cycle profiling showed strong G1 arrest and no difference in cell cycle state between control and drug treated cells after 24 hours (h) (Supp. Fig. 1a). Combretastatin A-4 (CA4), which binds tubulin at the colchicine site, was chosen as a reference destabilizer, and paclitaxel (PTX) as a reference stabilizer^18^ (Supp. Fig. 1b-e). We chose drug concentrations based on tubulin partitioning between polymer and dimer, and counting growing plus-tips labeled with end-binding protein 1 (EB1)^19^. Low doses (1nM CA4, 3nM PTX) were chosen to cause barely detectable effects, and high doses (100nM CA4, 300nM PTX) to cause complete loss of polymer (destabilizer) or complete loss of soluble dimer (stabilizer) (Supp. Fig. 1b-e). We then compared gene expression responses by RNA-sequencing (RNA-seq) of poly-adenylated (polyA+) mRNAs at 6 and 24h post-drug treatment. These data revealed differential expression of multiple genes under all conditions relative to basal expression (Benjamini-Hochberg corrected p-value<0.05, and >50 mRNA counts per million reads, Fig. 1a-d).

**Figure 1.**
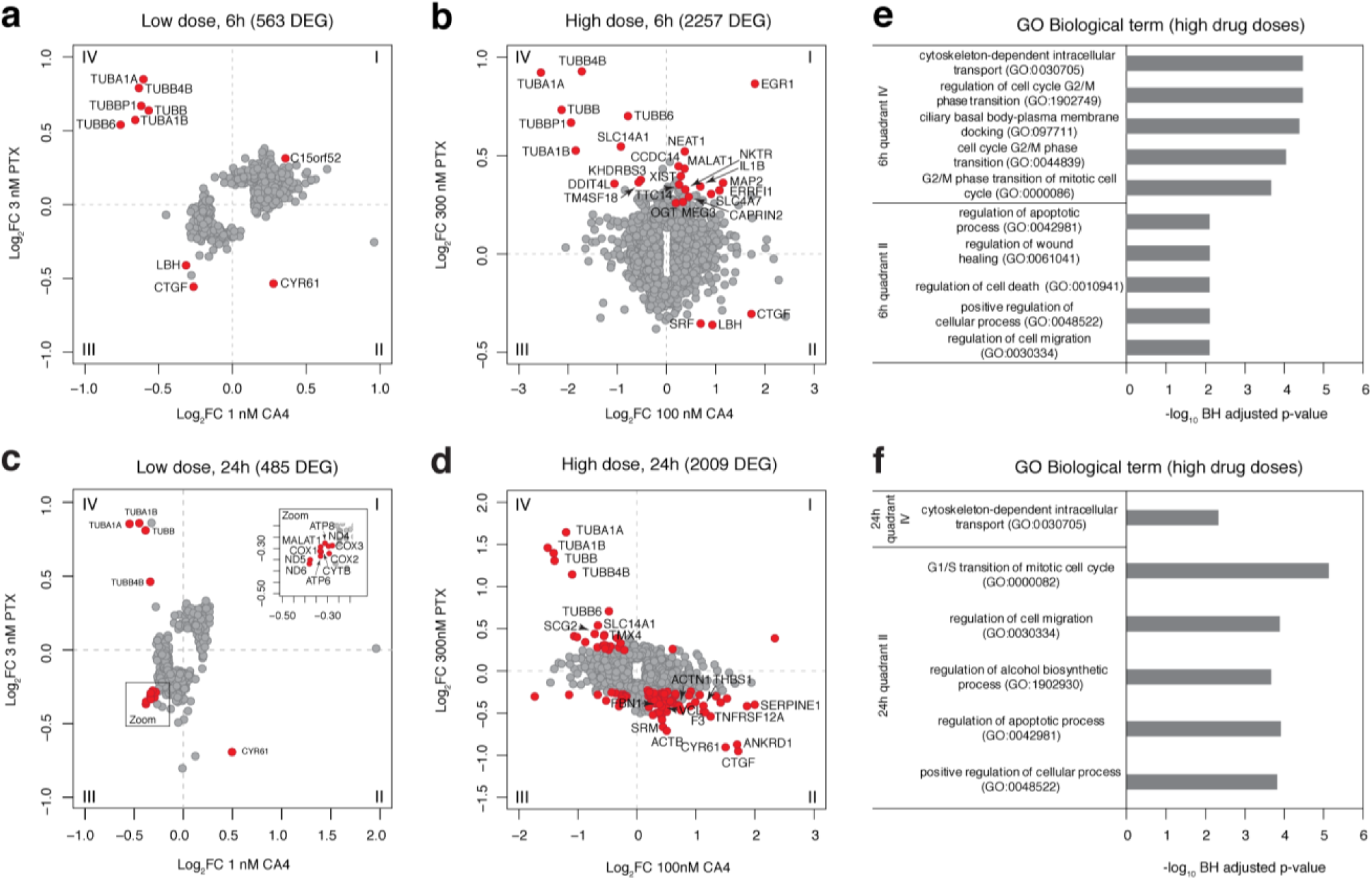
Microtubule damage triggers differential gene expression (DGE). **a-d)** DGE in RPE 1 hTert cells treated with microtubule poisons for 6 (a and b) or 24 h (c and d). Data are represented as average Log_2_ Fold Changes (Log_2_FC) from three independent biological replicates, relative to basal expression in DMSO-treated control cells. Plotted are expression profiles of genes that have p-value<0.05 and >50 mRNA reads per million, in 1 nM CA4 and 3 nM PTX-treated cells (a and c), and 100 nM CA4 and 300 nM PTX-treated cells (b and d). In red are depicted genes with false discovery rate (FDR) <0.02 for low drug doses, and <0.001 for high drug doses, of which names are printed for top 20 genes. Plot quadrants are labeled from I-IV. **e and f)** Gene set enrichment analysis (GSEA) based on expression profiles of genes from quadrants II and IV, from cells treated with high drug doses for 6 (b), or 24 h (d). Plotted are -Log_10_ Benjamini-Hochberg (BH) adjusted p-values of top five enriched gene ontology (GO) terms in category Biological Process, with BH-adjusted p-value <0.01.

Differential gene expression (DGE) relative to vehicle treated cells (Fig. 1a-d) is presented to reveal genes which respond similarly to both drugs (quadrants I and III) versus differentially (quadrants II and IV, Fig. 1a-d). As expected, high drug doses caused extensive gene expression changes. Surprisingly, sub-threshold doses also caused many significant changes, showing that quiescent cells can detect even mild perturbation of microtubule dynamics. Based on high drug dose expression profiles, we performed gene set enrichment analysis (GSEA), finding significantly enriched gene ontogeny (GO) terms (Fig. 1e-f). The most prominently enriched are genes involved in G1/S transition of the mitotic cell cycle, which were up-regulated in CA4-treated and down-regulated PTX-treated cells (Fig. 1f). These data are consistent with previous reports that microtubule destabilization promotes DNA replication and cell cycle re-entry, while stabilization inhibits it^20-22^. We further found highly enriched genes involved in apoptosis, wound healing, and cell migration, which were also upregulated upon microtubule destabilization and downregulated upon stabilization (Fig. 1e-f). Surprisingly, we found only one cluster of microtubule-related genes involved in cytoskeleton based-transport (6h and 24h post-treatment) that were differentially regulated by microtubule stabilization versus destabilization (Fig. 1e-f). We conclude that cells mount both differential and common responses to microtubule-stabilization versus destabilization. Furthermore, they clearly detect damage that is sub-threshold by biochemical tubulin-partitioning criteria. Notably, we did not observe changes in mitosis-related genes in quadrants I and III, consistent with the lack of mitotic arrest in our quiescent cultures. Some common DGE signatures were suggestive of general stress responses, such as apoptosis, and these deserve further analysis.

Tubulin genes stood out as most differentially regulated genes in quadrant IV in all four plots, particularly in the low dose perturbations (Fig. 1a-d). This observation was statistically confirmed by GSEA analysis, where tubulin differential expression drove enrichment of GO terms such as “cytoskeleton dependent intracellular transport” (Fig. 1e-f). We conclude that coordinated change in multiple tubulin mRNAs was the strongest response to drug-induced microtubule damage, with destabilizing drugs decreasing tubulin mRNA concentrations, and stabilizing drugs increasing them.

### Differential regulation of most tubulin isoforms upon MT damage

To test if all tubulin mRNAs undergo similar regulation, we extracted the expression profiles of all detected tubulin genes in our data set, and their total mRNA counts (Fig. 2a). We observed differential regulation of all highly expressed α- (TUBA) and β-tubulin (TUBB) mRNAs, but not centrosomal δ- (TUBD1) and ε-tubulins (TUBE1) (Fig. 2a). Moreover, we found clear differential expression of centrosomal γ-tubulin isoform 1 (TUBG1), and borderline regulation of isoform 2 (TUBG2) (Fig. 2a).

**Figure 2.**
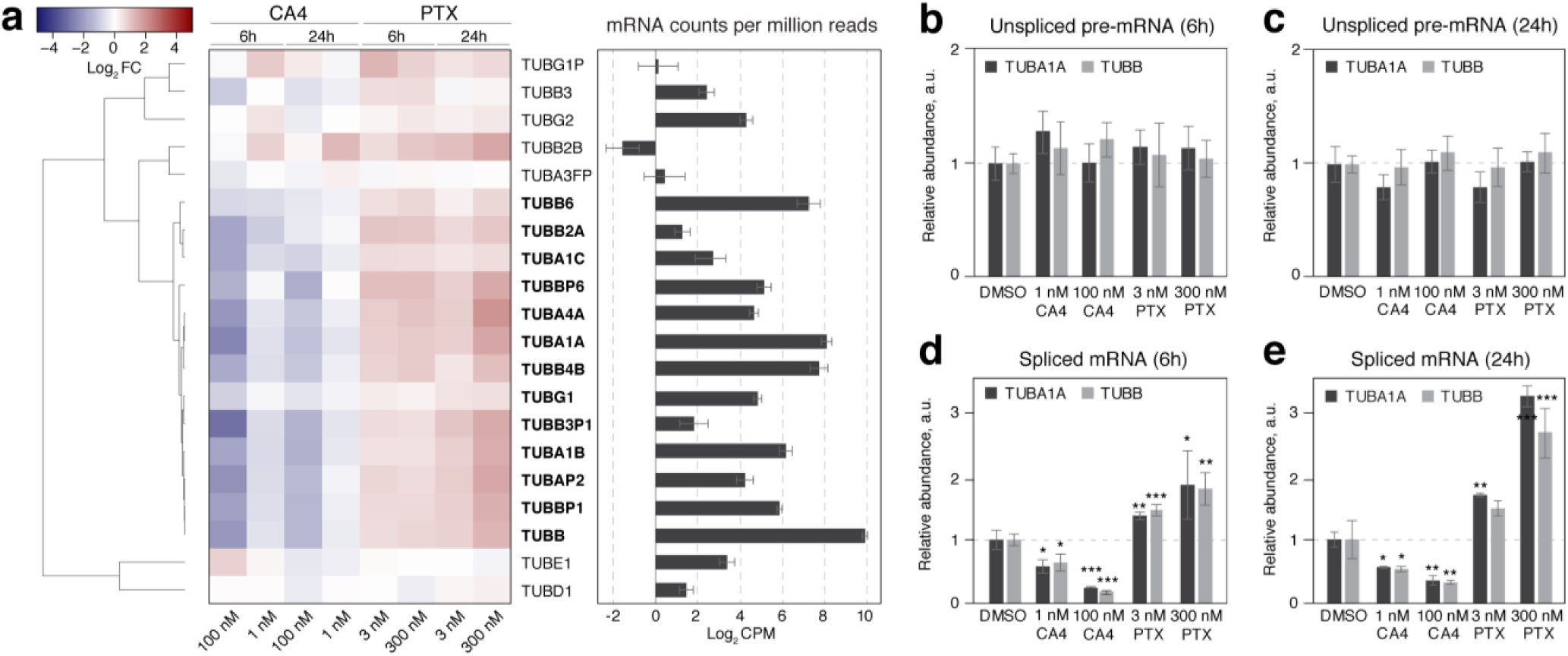
Microtubule damage triggers differential expression of all α- and β-tubulin isoforms. **a - left panel)** Expression profiles of all detected α- and β-tubulin isoforms in our DGE dataset. Dendrogram on the left represents Pearson distance between expression profiles. Each column of the heatmap represents differential gene expression in one treatment, labeled on x-axis above, and bellow the heatmap, relative to DMSO control. Each row represents a gene, labeled on y-axis. In bold are differentially expressed genes that have p-value <0.05. Color key is depicted in upper left corner. Data are represented as Log2FC relative to DMSO control. **a - right panel)** Average log_2_ mRNA counts for each gene per million detected reads (CPM) in DMSO-treated control samples are presented as a bar chart. **b and c)** Relative abundance of unspliced TUBA1A and TUBB pre-mRNA in control (DMSO), and cells treated with microtubule poisons (x-axis) for 6 (b) or 24 h (c). **d and e)** Relative abundance of spliced TUBA1A and TUBB mRNA in control (DMSO), and cells treated with microtubule poisons (x-axis) for 6 (d) or 24 h (e). All the expression profiles are normalized to a reference gene (GAPDH or RPL19), and to DMSO control. Error bars in all panels represent standard deviation from three independent biological replicates. * p-value<0.05, ** p-value<0.01, *** p-value<0.001 in paired Student T-test compared to DMSO control.

To generalize this finding, we re-analyzed two large, high-quality datasets deposited in Gene Expression Omnibus (GEO) database that profiled DGE response to microtubule damage. In an extensive study that compared PTX with eribulin (ERB, a microtubule destabilizer) treatment of many breast, ovarian, and endometrial cancer cell lines^15^, we confirmed differential regulation of all expressed α- and β-tubulins, and γ-tubulin isoform 1, (Supp. Fig. 2a). Importantly, re-analyzing a study that compared the effect of microtubule destabilizers colchicine, vinblastine, and vincristine on rat heart endothelial cells^23^, we show for the first time differential regulation of tubulin genes *in vivo*, (GEO GSE19290, Supp. Fig. 2b). We conclude that cells differentially regulate all the expressed α- and β-tubulin isoforms, and γ-tubulin isoform 1 upon microtubule damage, *ex vivo* and *in vivo*.

The microtubule damage-induced changes in tubulin mRNA concentrations that we observed were strongly suggestive of tubulin autoregulation, a post-translational gene-expression regulation mechanism^24^. RNA-seq of polyA+ mRNA does not distinguish between transcriptional and post-transcriptional regulatory mechanisms. To make this determination we established a reverse transcription quantitative PCR-based assay (RT-qPCR), to specifically measure transcriptional regulation through the expression levels of unspliced pre-mRNA, and post-transcriptional regulation through the expression levels of spliced mRNA (Supp. Fig. 2c).

Using this approach, we measured two highly-expressed tubulin genes, α-tubulin 1A (TUBA1A), and β-tubulin B (TUBB), and two control housekeeping genes, Glyceraldehyde 3-phosphate dehydrogenase (GAPDH), and ribosomal protein L19 (RPL19). We found no significant change in unspliced TUBA1A and TUBB pre-mRNA concentration in cells treated with CA4 or PTX (Fig. 2b-c), showing that microtubule damage did not change the rate of tubulin gene transcription. However, levels of mature, spliced TUBA1A and TUBB mRNAs significantly diminished in CA4-treated cells and increased in PTX-treated cells (Fig. 2d and e), consistent with our RNA-seq data. We conclude that post-transcriptional regulation of tubulin mRNA stability is the most prominent gene expression response to microtubule damage. Importantly, we did not observe co-regulation of any microtubule interacting proteins, such as microtubule associated, motor or plus-tip binding proteins. Thus, altered stability of microtubules only regulates the expression of tubulins, but not the other components of functional microtubules.

### Bioinformatic analysis of the autoregulation signature revealed new microtubule biology

We next sought to investigate if tubulin DGE is a general response to altered microtubule dynamics in conditions other than microtubule-targeted poisoning. The differential tubulin gene expression triggered by microtubule damage comprises a strong and specific signature that can be used to query publicly available DGE datasets in an unbiased manner, and with the expectation of finding novel conditions that regulate microtubules. To test this approach, we used CLustering by Inferred Co-expression^25^ (CLIC, www.gene-clic.org, Fig. 3a)—a bioinformatic tool that mines ∼3500 publicly available human and mouse RNA-profiling studies deposited in the GEO database. Importantly, most of these studies are not designed to research cellular response to microtubule damage, providing an unbiased approach that can potentially reveal new microtubule biology.

**Figure 3.**
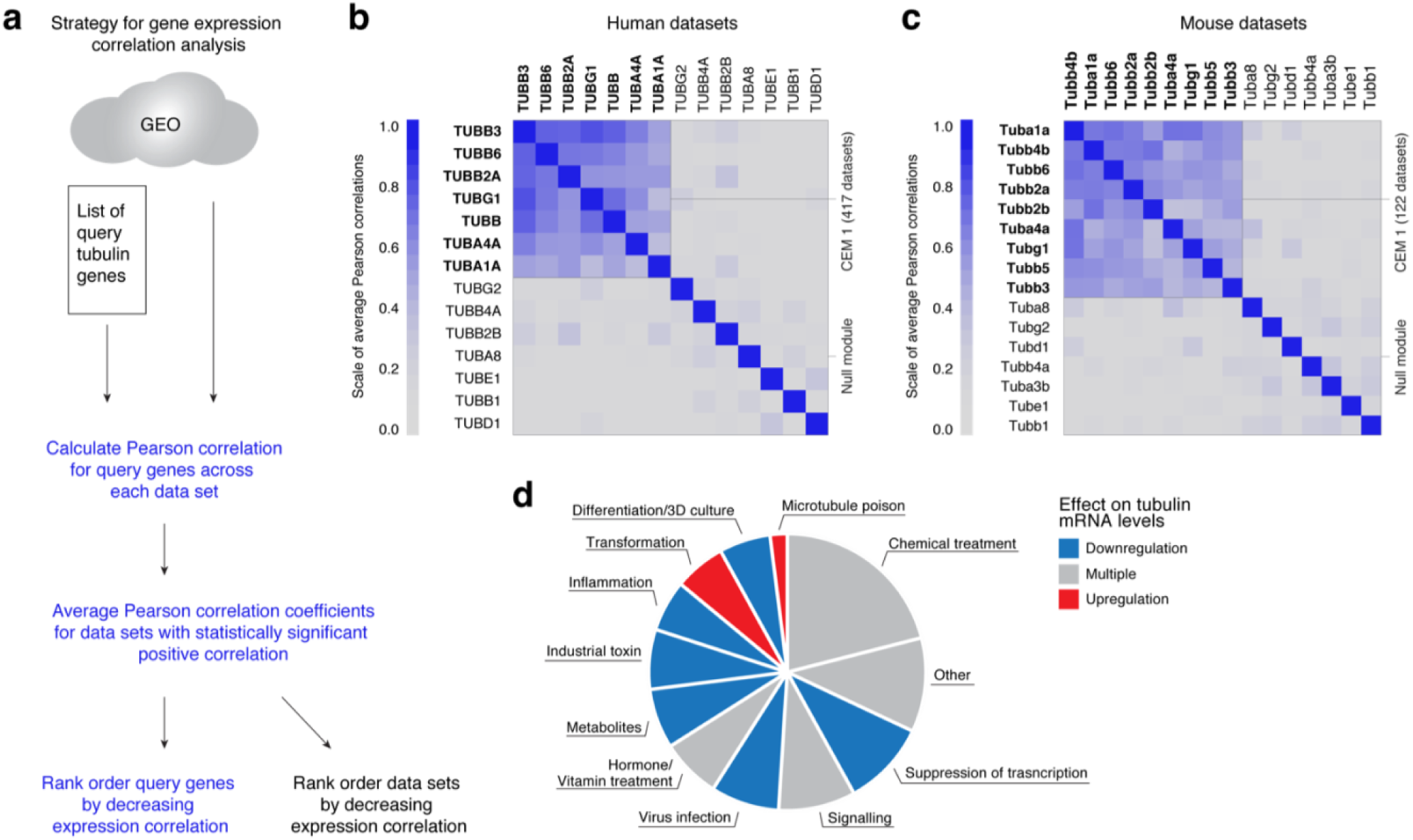
Cells coordinate expression of α- and β-tubulin isoforms. **a)** Scheme of the bioinformatic approach. **b and c)** Pearson expression correlation coefficients for a subset of abundantly expressed α- and β-tubulin isoforms, across 417 human (b), and 122 mouse (c) publicly available Affymetrix chip data sets. In dark red are genes that show strong expression correlation (Pearson correlation coefficient=1), and in gray are genes that show no expression correlation (Pearson correlation coefficient=0). Color keys are represented to the left of each heatmap. High expression correlation for a subset of tubulin genes is marked as cluster CEM1. Tubulin genes that do not correlate in expression are marked as Null module. **d)** Frequency diagram of groups of perturbations amongst the top 100 data sets with the highest tubulin gene expression correlation from the human platform.

We first investigated whether tubulin genes were coordinately expressed across many sample and perturbation types. We found high Pearson expression correlation coefficients for the most broadly expressed tubulin genes in 417 human (Fig. 3b), and 122 mouse studies (Fig. 3c). More specialized, and low abundance tubulin isoforms displayed lower expression correlation coefficients. The obtained expression correlations confirm that cells co-regulate the mRNAs for multiple α- and β-, and γ-tubulin isoform 1 in many cell and tissue types across a total of 539 different perturbations. These perturbations are candidates for novel conditions that regulate the microtubule cytoskeleton.

To understand what kinds of experimental perturbations induce co-regulation of tubulin genes, and potentially regulate the microtubule cytoskeleton, we rank ordered the human datasets by descending Pearson correlation coefficient (Supp. Table 2) and manually annotated the top 100 (Fig. 3d, Table 1). Amongst high-ranked studies were multiple investigations of established microtubule drugs, such as PTX, or toxins known to perturb microtubules, such as bisphenol-A (Table 1). These served as positive controls that our bioinformatic analysis returned studies where microtubules were perturbed. We also discovered many novel conditions, including virus infection, metabolite deprivation, exposure to industrial toxins, inflammation, and cell differentiation, which all tended to cause coordinated downregulation of tubulin mRNAs (Figure 3d, Table 1). Surprisingly, only one process, oncogenic transformation, consistently upregulated tubulin mRNAs (Figure 3d, Table 1). Hormone, vitamin, and chemical treatment, as well as perturbations of signaling, had molecule-specific effects on tubulin gene expression (Figure 3d, Table 1). In some studies, inspection of the CLIC-report showed that co-regulation of tubulin was not induced by the experimental perturbation in the title of the study, but rather by some other controlled and annotated experimental variable. For example, Steroid Receptor Coactivator-1 (SRC1) RNAi in A549 lung cancer cells did not change tubulin gene expression compared to control RNAi, but glucose withdrawal, which was included as a variable in the same study suppressed it^26^ (Supp. Fig. 3a).

**Table 1.**
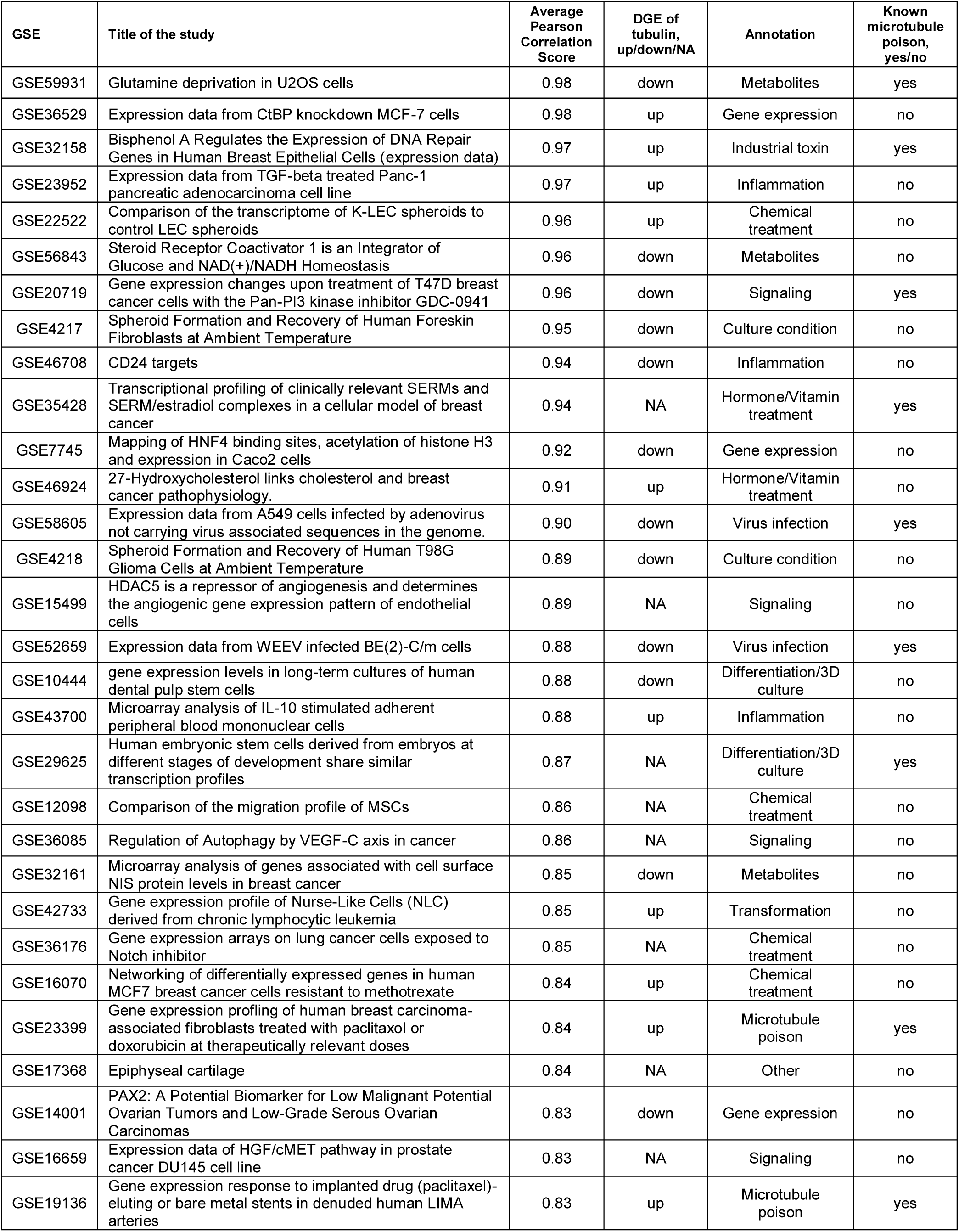
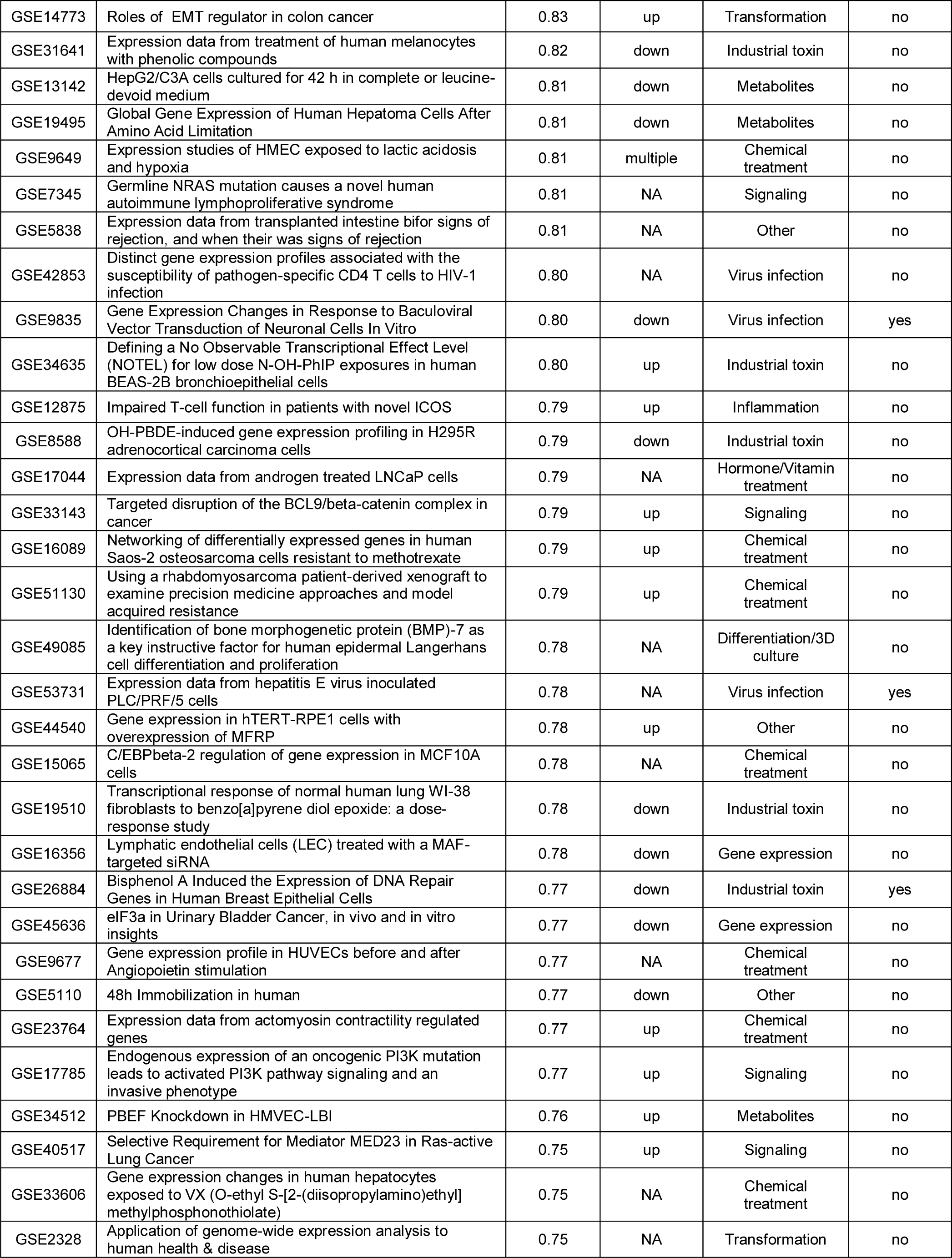
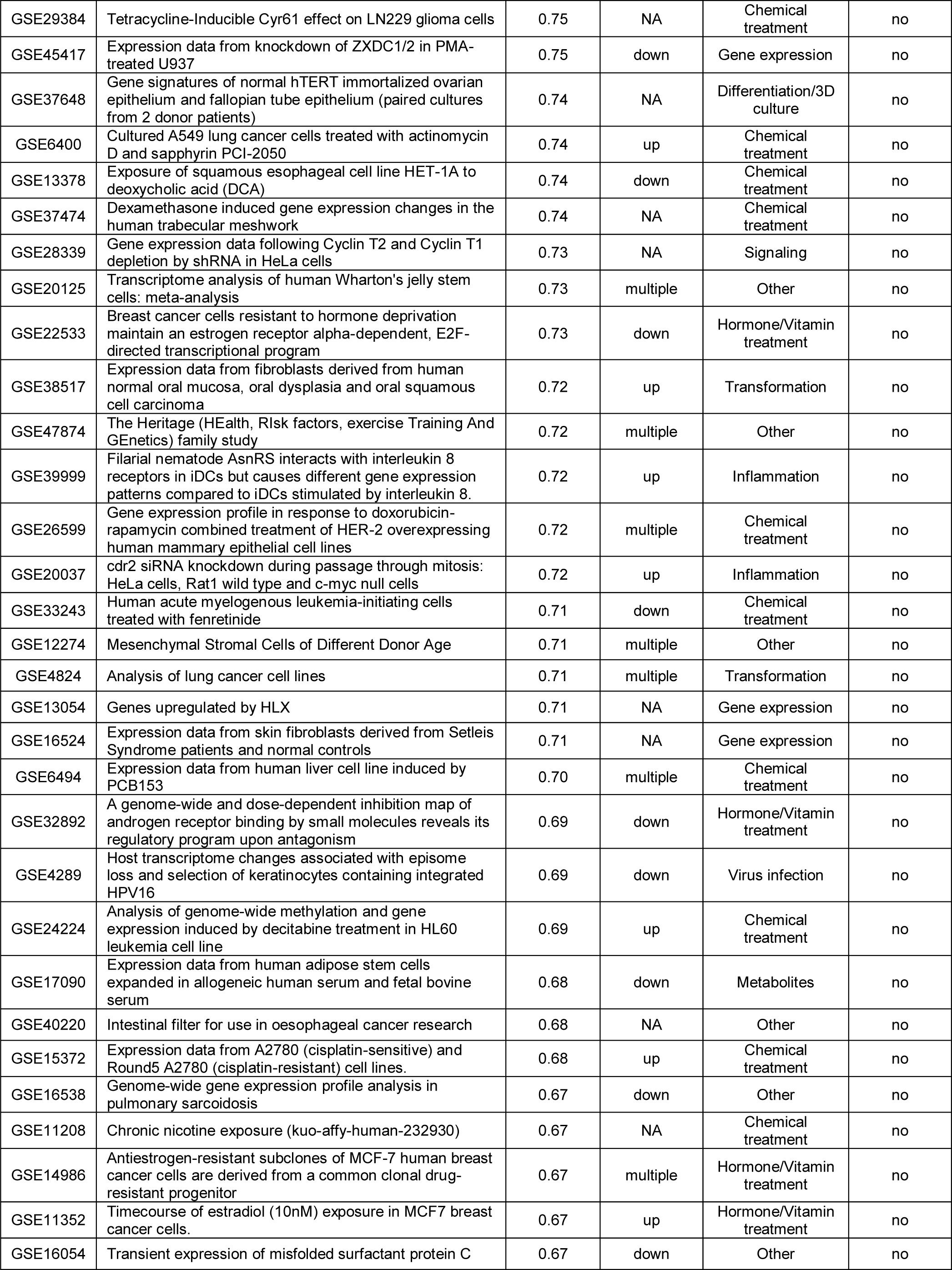
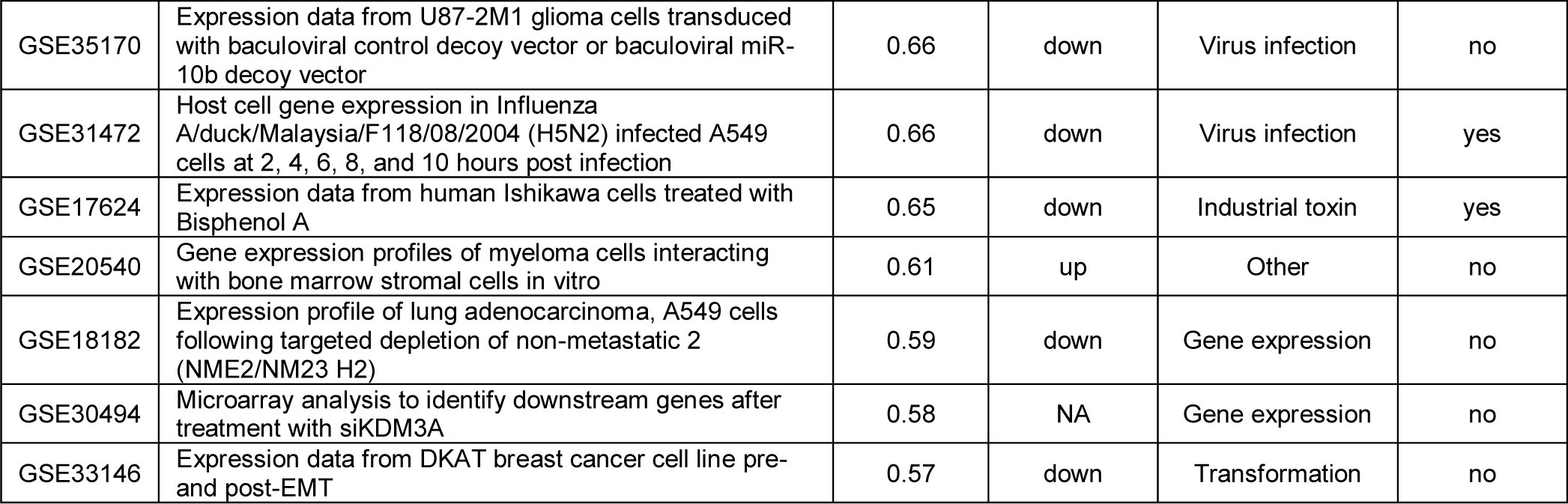
The top 100 datasets with highest tubulin gene expression Pearson correlation from the human platform. Annotated are studies rank ordered by decreasing tubulin gene expression Pearson correlation, with their associated GEO identifiers. Extracted from the CLIC-report are differential tubulin gene expression. Unless not detected (NA), downregulated (down), or upregulated (up) tubulin mRNA levels are annotated. Based on published literature, perturbations in the listed studies are annotated into 11 groups, and their effect on microtubule cytoskeleton is marked as known (yes), or unknown (no).

To investigate whether our bioinformatic analysis returned new microtubule-biology, we selected several perturbations, starting with nutrient deprivation, for validation by RT-qPCR, and subsequent biochemical and microscopy-based inspection of the microtubule cytoskeleton. We confirmed that glucose and glutamine deprivation lower tubulin mRNA levels in cancer cells lines, further finding that this occurred by a mixture of transcriptional and mRNA stability mechanisms (see Supp. Fig. 3 and related text). In both scenarios, nutrient deprivation caused destabilization of the microtubule cytoskeleton (Supp. Fig. 3d and h). We conclude that various physiological and damaging inputs change tubulin gene expression either transcriptionally, thus affecting the microtubule cytoskeleton, or post-transcriptionally through changes in microtubule dynamics.

### PI3K activity increases tubulin gene expression via microtubule stability and autoregulation

Multiple investigations of PI3K signaling scored highly in our bioinformatics search, including studies where genetics were used to up-regulate the pathway activity, and small molecule inhibitors to down-regulated it. In each case, PI3K pathway activation correlated with an increase in tubulin mRNA levels (GEO GSE17785), and pathway inhibition with a decrease^27^ (Table 1, Supp. Table 1). One CLIC-report suggested that MCF10A cells that expressed constitutively active PI3K with a mutation in the kinase domain^28^ (H1047R) had increased tubulin mRNA levels (GEO GSE17785, Fig. 4a). PI3K inhibition with GDC-0941 prevented this upregulation of tubulin mRNA levels, suggesting involvement of PI3K in concerted regulation of tubulin expression (Fig. 4a). Using RT-qPCR, we found that the transcription rate of TUBA1A and TUBB in H1047R cells was unchanged, compared to parental cell line (Fig. 4b). Levels of spliced TUBA1A and TUBB mRNAs (Fig. 4c), however, were significantly higher in H1047R cells, suggesting post-transcriptional regulation of tubulin mRNA. To directly test if PI3K was involved in the regulation of tubulin mRNA stability, we measured tubulin mRNA levels in parental and H1047R-mutant cells treated with the PI3K inhibitor GDC-0941^27^. PI3K inhibition in parental cell line increased transcription, but not the stability of tubulin mRNAs (Fig. 4b-c). Importantly, while PI3K inhibition in H1047R-mutant cell line increased the transcription of TUBA1A, but not TUBB, it restored normal levels of spliced TUBA1A and TUBB mRNA, comparable to parental cell line (Fig. 4b and c). To test if these results were more general, we measured TUBA1A and TUBB gene expression in another cell line carrying an activating mutation in the helical domain of PI3K (E545K^28^). We found the rate of TUBA1A and TUBB transcription unchanged (Fig. 4b), while the levels of spliced TUBA1A and TUBB mRNA were significantly higher in E545K compared to parental cells (Fig. 4c), again consistent with regulation by mRNA stability. Treatment of E545K cells with GDC-0941 PI3K inhibitor restored normal levels of spliced TUBA1A and TUBB mRNA (Fig. 4c), suggesting that PI3K activity regulated tubulin mRNA levels. As in parental and H1047R cells, transcription of TUBA1A and TUBB was higher in E545K-mutant cells treated with GDC-0941 compared to control-treated cells, suggesting transcriptional activation triggered by PI3K inhibition (Fig. 4b). We conclude that, despite activating transcription, PI3K inhibition post-transcriptionally suppresses tubulin mRNA levels, presumably through autoregulation.

**Figure 4.**
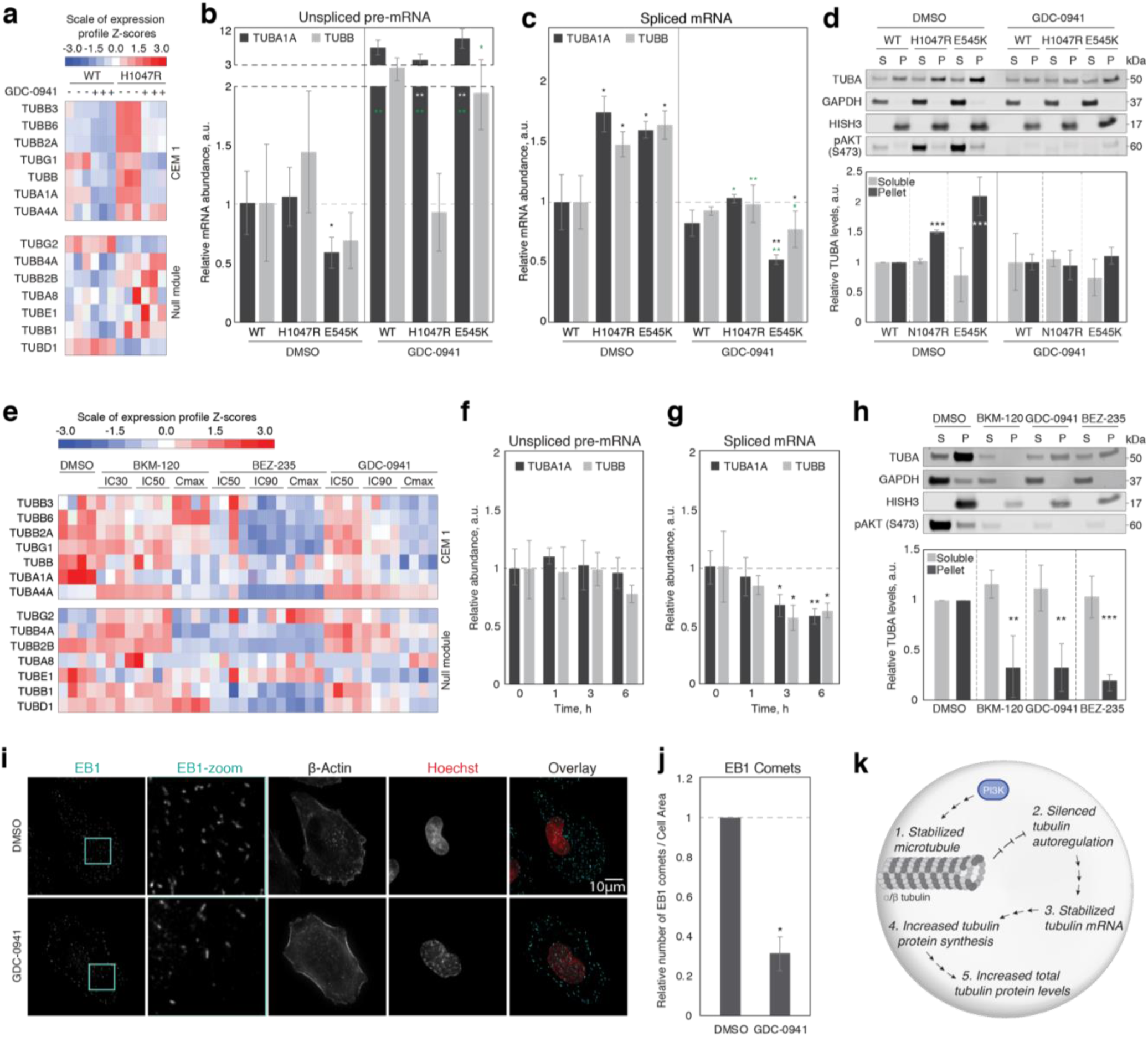
PI3K activity increases tubulin gene expression through autoregulation. **a)** DGE in parental and cells expressing constitutively active PI3K H1047R-mutant cells, treated with DMSO or indicated PI3K inhibitors for 4 h. **b and c)** Relative expression of TUBA1A and TUBB unspliced pre-mRNA (b), and spliced TUBA1A and TUBB mRNA (c) in parental, and PI3K mutant cell lines H1047R, and E545K, treated with DMSO control or 1 μM GDC-0941 for 4h. All the relative gene expression data are normalized to housekeeping gene GAPDH or RPL19, and to DMSO-treated parental cells. **d)** Tubulin partitioning to unpolymerized (soluble, S) and polymerized (P) normalized to loading control (GAPDH for S; HISH3 for P) and to DMSO-treated parental cells, in parental and cells expressing constitutively active PI3K mutants H1047R and E545K, treated with DMSO or 1 μM GDC-0945 for 4 h. **e)** DGE in A2058 cells treated with DMSO control or the indicated concentrations of PI3K inhibitors, for 6 h (GEO series GSE66343^27^). **f-g)** RT-qPCR in A2058 cells treated with 1 μM GDC-0941 for indicated periods of time (x-axes). **h)** Tubulin partitioning to unpolymerized (S) and polymerized (P) normalized to loading control (GAPDH for S; HISH3 for P), and to DMSO-treated cells, in control and cells treated for 6 h with indicated 1 μM PI3K inhibitors. **i)** Representative immunofluorescence images of control and cells treated with 1 μM GDC-0941 for 6h, and stained with anti-EB1 antibody (cyan), anti-β-actin, and Hoechst (red). **j)** Relative number of detected EB1 comets per cell area, in DMSO control and A2058 cells treated with 1μM GDC-0941 for 6 h, normalized to DMSO-treated cells (100-200 cells). **k)** A proposed model describes the mechanism through which PI3K signaling regulates tubulin gene expression through modification of tubulin autoregulation. Bar plots in all panels represent average values, and error bars standard deviations from three independent biological replicates. * p<0.05, ** p<0.01, *** p<0.001 in Paired Student T-test, in all panels. Black and white asterisk represent statistical significance relative to DMSO-treated parental cell line, green asterisks represent statistical significance relative to DMSO-treated control of the same cell line.

We next sought to determine if PI3K activity regulated the stability of tubulin mRNAs via changes in microtubule stability and altered ratio of soluble tubulin dimer and polymer. To test if PI3K activation causes a decrease in the concentration of soluble tubulin dimer, we performed biochemical tubulin partitioning on parental, H1047R and E545K cells. We found that both H1047R and E545K cell lines had less soluble tubulin dimer and more microtubule polymer than the parental cell line (Fig. 4d). PI3K inhibition in these cells increased levels of soluble tubulin dimer and decreased microtubule polymer, comparable to the levels observed in parental cells (Fig. 4d). These data are consistent with a model where PI3K activity increases microtubule stability, leading to a decrease in soluble tubulin and an increase in tubulin mRNA stability via the autoregulation pathway. PI3K inhibition with a small molecule reverses these changes. While these data do not reveal the mechanism by which PI3K regulates microtubule stability, they are consistent with published models^29^.

To generalize our findings, we investigated a second cell line, A2058, where our bioinformatic analysis revealed down-regulation of tubulin mRNAs upon PI3K inhibition. PI3K is constitutively active in these cells (Figure 4h). We treated A2058 cells with the PI3K inhibitors BKM-120^30^ (inhibitor for p110α/β/δ/γ), BEZ-235^31^ (a dual PI3K and mTOR inhibitor for p110α/γ/δ/β), or GDC-0941^32^ (inhibitor for PI3Kα/δ). In each case drug treatment decreased the levels of tubulin mRNA (Fig. 4e). Performing RT-qPCR on cells treated with 1 μM GDC-0941 (IC90^27^), we found that PI3K inhibition did not affect transcription of tubulin genes (Fig. 4f), but it post-transcriptionally down-regulated tubulin mRNAs (Fig. 4g), consistent with regulation through mRNA stability. Biochemical partitioning of tubulin revealed that treatment with 1 μM BKM-120, BEZ-235, or GDC-0941 indeed caused an increase in levels of soluble tubulin dimer and a reduction of microtubule polymer (Fig. 4h). To further confirm microtubule destabilization upon PI3K inhibition, we used immunofluorescence-based microscopy. We fixed and stained control and PI3K-ihibited cells, using anti-EB1 antibody, and counted the number of EB1-positive microtubule plus-tips per cell area as a read-out for growing microtubules. Our data showed significant reduction in the number of growing microtubules in cells treated with 1 μM GDC-0941compared to control-treated cells (Fig. 4i and 4j).

Several studies have identified off-target destabilization of microtubules by kinase inhibitors, so it was important to test for this possible artifact. In RPE 1 hTert cells, with low basal PI3K activity, we did not observe changes in the number of EB1-positive microtubule plus-tips upon PI3K inhibition with GDC-0941 or BEZ-235, suggesting absence of off-target activity on tubulin (Supp. Fig. 4a-b). Consistent with previous reports of off-target effects on microtubules^33^, high dose of PI3K inhibitor BKM-120 caused a reduction in the number of EB1-positive microtubule plus-tips (Supp. Fig. 4a-b).

Taken together, we conclude that PI3K activity positively regulates tubulin levels in cancer cells via microtubule stabilization, which lowers the concentration of soluble tubulin dimer, thus increasing tubulin mRNA stability through the autoregulation pathway. Our findings show for the first time that tubulin autoregulation via mRNA stability mediates changes in total tubulin concentration triggered by an important signaling pathway. PI3K inhibition in cell lines with low constitutive kinase activity did not perturb microtubules (Fig. 4d, Supp. Fig. 4a-b) suggesting that this regulation is selective for cancer cells, and normal physiological states where PI3K is strongly activated.

## Conclusions

Our data show that quiescent cells can detect both subtle and strong perturbation of microtubule dynamics and mount robust gene expression responses. Moreover, our study is the first to tease apart the opposite effects of microtubule-destabilization versus stabilization on gene expression from their similar effects on mitotic arrest. Our findings begin to validate the idea that an interphase “Microtubule Integrity Response” (MIR) pathway exists. Tubulin genes were the most responsive, especially to mild microtubule perturbation. Our RT-qPCR data confirm that this regulation involves a previously reported pathway, tubulin autoregulation, where control occurs at the level of mRNA stability, not transcription. The molecular mechanism of tubulin autoregulation is not known, but it is thought to involve co-translational degradation of tubulin mRNAs by a pathway that is sensitive to the concentration to soluble tubulin dimer^13,14^.

Due to limitations in available technology at the time of discovery, early studies of tubulin auto-regulation could only measure total α- and β-tubulin mRNA regulation, and not the regulation of specific isoforms^12-14^. Our study is the first to show that autoregulation extends to all the expressed α- and β-tubulin genes. Understanding the molecular mechanism and physiological relevance of tubulin autoregulation will be important for the discovery and characterization of the hypothetical MIR.

Prior work on tubulin autoregulation was confined to microtubule-drug induced effects in a few cell types in culture^13,34,35^. Using bioinformatics to query public datasets, we provide evidence that co-regulation of expressed α, β and γ-tubulins is observed in response to stimuli other than microtubule-targeting drugs, in tissues, and in cycling and quiescent cell cultures. Moreover, we show for the first time that this pathway is part of concerted cellular responses to many different stimuli. In the case of PI3K signaling, we find that activation of the kinase stabilizes microtubules, thus decreasing the pool of soluble tubulin dimer and stabilizing tubulin mRNA. This is consistent with the proposed mechanism of tubulin autoregulation. Our findings, therefore, demonstrate the ubiquity and importance of autoregulation in controlling tubulin gene expression, and very likely microtubule biology in general.

### Autoregulation suggests functions of tubulin beyond building microtubules

A striking feature of our DGE data in quiescent RPE1 hTert cells, which was also seen in two large GEO studies that we re-analyzed, is a lack of co-regulation of other microtubule components with tubulin: tubulin autoregulation signature extends to all expressed tubulin, but not to microtubule-associated, plus-tip-binding, or motor proteins, nor are these components co-regulated on the transcriptional level in response to microtubule damage. This finding is quite different from coordinated gene expression in other biological responses, for example many of the genes required to build lysosomes^36^, or mitochondria^37^ are coordinately regulated. The specific regulation of all expressed tubulins, but not other microtubule proteins, suggests that cells care more about the concentration of tubulin than other components of the microtubule cytoskeleton. Perhaps, this simply reflects the central role of tubulin in building microtubules. A related puzzle is that autoregulation is counter-homeostatic from the perspective of microtubule mass, at least in response to microtubule drug perturbation. CA4-treatment reduces microtubule mass, yet cells respond by destabilizing tubulin mRNA, and reducing protein levels, which would seem to exacerbate the problem, not correct it. The converse is true for PTX. An intriguing possible explanation for both the apparent paradoxes is that soluble tubulin dimer has some function other than building microtubules, which necessitates tight control of its concentration. For example, soluble tubulin was proposed to gate Voltage-Dependent Anion Channels (VDAC) in mitochondria^38^, and it is conceivable that autoregulation evolved to regulate this function, not microtubule assembly. Another possibility is that autoregulation evolved to balance the synthesis of α- and β-tubulin subunits, in which case other components of the microtubule cytoskeleton are irrelevant. But if this is the case, why is TUBG1 also co-regulated? These are speculations, but they point out that we truly do not understand the adaptive benefit of tubulin autoregulation, despite its frequent occurrence in cellular physiology, as revealed by our bioinformatics analysis.

### Discovery of physiological processes that change microtubule stability from genomic data

Research in the microtubule field has been driven largely by microscopy and biochemistry, and genomic approaches have been neglected. Our CLIC search shows that bioinformatic mining of public datasets can be used as a starting point for discovery. As a positive control, this approach returned multiple studies of microtubule perturbing agents, such as PTX and bisphenol-A. Unexpectedly, we uncovered many more previously unreported conditions that perturb tubulin gene expression, which remain to be investigated.

Our mechanistic follow-up study showed that PI3K activity promotes stabilization of tubulin mRNA. This effect is achieved through microtubule stabilizing activity of PI3K signaling and tubulin autoregulation. Notably, the effect of autoregulation on basal tubulin levels, and on response to PI3K inhibitors, was stronger in cells carrying an activating mutation in PI3K. Activating mutations in PI3K cause many changes in growth and metabolism in cancer cells^39^. Our data add change in tubulin levels to that picture, which could be significant for successful mitosis, response to microtubule drugs, and metastasis pathways regulated by microtubules.

Our genome-scale data analysis, and PI3K investigation point to the importance of tubulin autoregulation in normal biology and cancer pathophysiology. Tubulin autoregulation was a central concern in the microtubule field in the 1980s, but it was abandoned as the field moved towards biophysical directions in the 2000s. We feel it is now important to investigate the full molecular basis of the autoregulation pathway, and extend the pioneering work on the role of the N-terminal peptide of nascent β-tubulin^14^.

## Materials and methods

### Cell culture and drug treatments

All the cell lines used in this study were grown at 37 °C with 5 % CO_2_ in a humidified incubator. hTert-RPE1-eGFP-EB3 (a kind gift from D. Pellman, HMS, Boston, USA), and hTert-RPE1 (ATCC, Manassas, VA) cell lines were grown in Dulbecco’s modified medium (nutrient mixture F12, DMEM/F12) supplemented with 10 % FBS, and 1 %(vol/vol) penicillin/streptomycin (pen/strep). Parental and PI3 kinase mutant MCF10A cell lines, H1047R and E545K (a gift from J. Brugge, HMS, Boston, USA), were grown as previously described (ref). Prior to drug treatment, Parental and PI3 kinase mutant MCF10A cell lines, H1047R and E545K cell lines were washed 2 times with 1x PBS and grown in medium lacking EGR^28^. Cells were incubated overnight. A2058 cells (kind gift from P. Sorger, HMS, Boston, USA), U2OS, and A549 cells (ATCC, Manassas, VA) were grown in DMEM supplemented with 10 % FBS and 1 % (vol/vol) pen/strep. Microtubule drugs were used at 1 nM or 100 nM for CA4, or 3 nM or 300 nM for PTX, for 6 and 24 hours. PI3K inhibitors BKM-120, BEZ-235, and GDC-0941 (a gift from Nathanael Gray, HMS, Boston, USA) were administered at 1 μM. All the microtubule poisons and PI3K inhibitors were dissolved in DMSO, and 0.01% DMSO was used as vehicle control. D-glucose free medium was prepared using D-glucose free DMEM (Thermo-Fisher Scientific, USA) supplemented with 10 % dialyzed FBS (Thermo-Fisher Scientific, USA), and 1 % vol/vol pen/strep (Thermo-Fisher Scientific, USA). Media with D-glucose was prepared by adding D-glucose (Sigma Aldrich, USA) to the glucose-free medium at 4.5 g/l final concentration. L-glutamine free medium was prepared using L-glutamine and sodium-pyruvate free DMEM (Corning, USA) supplemented with 10 % dialyzed FBS, 1 % sodium-pyruvate (Corning, USA), and 1 % vol/vol Pen/Strep. Media with L-glutamine was prepared by adding L-glutamine (Corning, USA) to the L-glutamine-free medium at 2 mM final concentration. For D-glucose and L-glutamine deprivation, standard growth medium was removed, cells were washed 3 times with 1x PBS, and media with (control) or without D-glucose or L-glutamine was added to cells.

### Flow cytometry cell cycle profiling

In the flow cytometry validation experiments, RPE 1 hTert cells were grown per treatment condition on 10 cm petri-dishes for 1 day for cycling culture, and 5 days after reaching full confluency for quiescent cultures. Cells were then treated with DMSO control, or 100 nM CA4, or 300 nM paclitaxel PTX for 6h. For each condition cells were detached from the dish in 0.05 % Trypsin solution, resuspended in 1x PBS supplemented with 1 % FBS and spun down by centrifugation (400 g for 5 min), followed by dispersion of the pellet into a single-cell suspension in 1X PBS. Cells were then fixed in 4 % PFA for 20 min on ice, stained with DAPI (Invitrogen, USA, #D1306), and analyzed on a BD LSRII flow cytometer (BD Biosciences, USA). The dye was excited with a 405 nm laser and emitted fluorescence was detected with a 450/50 bandpass filter. For data analysis, FlowJo v.X.0.7, and the built-in cell cycle quantification platform were used (the univariate model without any adjustments).

### RNA Sequencing and data analysis

Total RNA was collected and purified using the PureLink RNA Mini Kit (Invitrogen, Thermo Fisher Scientific, USA). RNA concentration and quality were determined using NanoDrop and Bioanalyzer respectively, and 500 ng of purified RNA was used as input for the Illumina TruSeq Stranded mRNA Library Prep Kit (Illumina, USA). Barcoded libraries were pooled and sequenced by The Bauer Core Facility at Harvard University, where the pooled library was quantitated using KAPA and single-end sequenced on an Illumina NextSeq (Illumina, USA). RNA-seq reads were mapped using STAR^40^ (version 2.1.0j) and processed using HTSeq-count^41^ (version 0.6.1). GRCh38 reference genome and transcript annotations were used for gene mapping; Entrez Gene identifiers and org.Hs.eg.db database were used for genome wide annotation^42^. Differential gene expression and statistical analysis were performed using edgeR package^43^. Genes with >50 reads per million and a fold change significantly different from zero in Wilcoxon signed-rank test (p< 0.05), were marked as differentially expressed genes, based on three biological replicates. EnrichR was used for gene set enrichment analysis^44,45^, and gene sets with Benjamini-Hochberg adjusted p-values p<0.01 were considered statistically significant.

### Bioinformatic analysis

We used CLIC-gene online tool for bioinformatic analysis of tubulin expression correlation^25^. Querying a list of all tubulin genes against human and mouse dataset platforms, the algorithm calculated average Pearson expression correlation for each dataset within one platform separately, and using expression correlation coefficients obtained from the top-ranked 417 human and 122 mouse differential gene expression data sets. The 417 human data sets with high tubulin expression correlation were then manually annotated.

### RT-qPCR

Reverse transcription was performed from 500ng of purified total RNA, using SuperScript IV (Invitrogen, USA) and random hexamer primers, according to manufacturer’s protocol. RT-qPCR reaction was performed using 5 ng of cDNA and 2x SYBRGreen master mix (Thermo Fisher Scientific, USA) on a BioRad thermocycler (BioRad, USA). For each reference and gene of interest, two sets of primers were designed: one set of primers amplified specifically unspliced pre-mRNA, while the other set of primers amplified specifically spliced mRNA. Primer sequences are listed in Supp. Table 1. All primer pairs were validated by PCR followed by gel-electrophoresis (data not shown). PCR products were further submitted to Sanger sequencing and subsequently mapped against the genome, confirming correct product amplification (data not shown). RT-qPCR data analysis was performed using the ddCt method^46^. Statistical significance was determined using two-tailed paired Student T-test and p<0.05.

### Western blotting and tubulin partitioning

Whole cell extracts for immunoblot analysis were prepared by cell lysis in 2x SDS sample buffer containing 50 mM Tris-HCl pH 6.8, 2 % SDS, 10 % glycerol, 1m% β-mercaptoethanol, 12.5 mmM EDTA, and 0.02 % bromophenol blue in water. Samples were denatured at 100 °C for 10 min, prior to SDS-PAGE gel electrophoresis on 1.5 mm NuPAGE Novex 4-12 % Bis-Tris Protein Gels (Thermo Fisher Scientific, USA), using Mini Trans-Blot Cell system (BioRad, USA).

For tubulin partitioning, cells were grown in 12-well dishes to 70-80 % confluence. Prior to lysis cells were washed 1x with 1xPBS pre-warmed to 37 °C, and PBS was aspirated. Unpolymerized tubulin was extracted in 300 μl of tubulin extraction buffer containing 60 mM PIPES (pH 6.8), 25 mM HEPES (pH 7.2), 10 mM EGTA (pH 7-8), 2 mM MgCl_2_, 0.5 % TritonX-100, 10 μM Paclitaxel (Sigma Aldrich, USA), and a protease inhibitor tablet (Roche, USA), for 2 min at room temperature. Extraction buffer with solubilized protein was collected and mixed with 100 μl 4x SDS sample buffer. The remaining material, containing polymerized tubulin, was then lysed with 400μl 1x SDS sample buffer. Samples were denatured at 100 °C for 10min. Prior to SDS-PAGE gel electrophoresis, equal aliquots of samples were concentrated 2x by evaporation.

For all immunoblots, Precision Plus Protein Dual Color Standard ladder was used (BioRad, USA). Dilutions and primary antibodies used: 1:15’000 GAPDH (Cell Signaling Technologies, USA, #2118), 1:10’000 alpha-tubulin (DM1alpha, Millipore, USA, #05-829), 1:10’000 Histone H3 (Cell Signaling Technologies, USA, #4499), and 1:500 phospho-Akt S437 (Cell Signaling Technologies, USA, #9271). The following secondary antibodies were used: 1:15’000 goat anti-mouse DyLight 680 conjugated (Thermo Fisher Scientific, USA, #35518), 1:15’000 goat anti-rabbit DyLight 800 conjugated (Thermo Fisher Scientific, USA, #35571). All the blots were imaged and analyzed using The Odyssey Infra-Red Imaging System (LI-COR, USA) equipped with Image Studio software. Statistical analysis and data plotting was performed in Microsoft Excel.

### Quantitative immunofluorescence

For quantitative immunofluorescence, cells were seeded and grown on coverslips overnight, prior to drug treatment. For immunostaining of EB1, cells were washed once with 1xPBS at 37 °C, and fixed in cold methanol at −20 °C for 20 min. Cells were then permeabilized in 4 % paraformaldehyde in PBS at room temperature for 20 min. After blocking in 2 % bovine serum albumin in PBS, primary antibodies were incubated at room temperatures for 1h. The following primary antibodies and dilutions were used: 1:500 mouse anti-EB1 (Cell Signaling, USA, #2164), 1:1’000 rabbit anti-β-actin (Cell Signaling, USA, #4970), 1:15’000 Hoechst 3342 (Thermo Fisher Scientific, USA, #H3570). Cells were then stained with secondary antibodies at room temperature for 30 minutes. The following secondary antibodies and dilutions were used: 1:400 Alexa Fluor Goat Anti-Rabbit IgG (Life Technologies, USA, #11008), and 1:400 Alexa Fluor Goat Anti-Mouse IgG (Life Technologies, USA, #11001). Three-dimensional image stacks of interphase cells were acquired in 0.2 μm steps using 60x NA 1.42 objective on an Olympus DeltaVision microscope (GE Healthcare, USA) equipped with DAPI/FITC/TRITC/CY5 filter set (Chroma, USA), and a sCMOS 5.5 camera (PCO, USA). EB1 comet counting and cell area measurements were performed on maximum-intensity z-stack projections, using a custom-made ImageJ macro, which is available upon request. Statistical analysis and data plotting were performed in R.

### Statistical analysis

For gene expression profiling, statistical analysis was supplied as part of edgeR package, and performed according to the user manual. For tubulin gene expression correlation, statistical analysis was performed using the online CLIC-gene tool. All the Student T-test analysis were performed considering two-tailed distribution, and two degrees of freedom.

## Acknowledgements

We are grateful to Kristina Holton and Research Computing Group at Harvard Medical School for their help with differential gene expression analysis, and Vamsi Mootha and Yang Li (Howard Hughes Medical Institute and Department of Molecular Biology, Massachusetts General Hospital, and Department of Systems Biology, Harvard Medical School, Boston, MA, USA) for help with the online CLIC-tool (www.clic-gene.com). We thank David Pellman, Joan S. Brugge, Nathanael Gray, Peter Sorger, Galit Lahav, and Marc Kirschner (Harvard Medical School, Boston, MA, USA) for sharing equipment, reagents, and cell lines with us. I.G. wishes to thank Luca Gerosa for help with RNA-seq data analysis and for inspiring discussions, Doaa Megahead for help with cell cycle profiling, and Javier J. Pineda for help with EB1 comet tracking (Department of Systems Biology, Harvard Medical School, Boston, MA). We thank Nikon Imaging Center, Flow Cytometry Facility, and Harvard Program in Therapeutic Science at Harvard Medical School, Boston, USA. We thank Sichen Shao (Cell Biology Department, Harvard Medical School, Boston, MA, USA) for discussions and critical reading of the manuscript, and the Mitchison lab members for valuable discussions. I.G. is a Merck Fellow of the Damon Runyon Cancer Research Foundation (DRG:2279-16). T.J.M. is currently supported by NIH-GM39565. S.A.B. is currently supported by NIH P50 GM107618.

## Author contributions

I.G. and T.J.M. conceived the project together. I.G. designed the experiments with T.J.M. All the experiments were performed by I.G. RNA-seq library was prepared by S.A.B. The manuscript was written by I.G. with input from S.A.B., and T.J.M.

