## Supplementary information for "Tubulin mRNA stability is sensitive to change in microtubule dynamics caused by multiple physiological and toxic cues"

**Supplementary figure legends**

**Supplementary Figure 1. Sub-threshold and saturating effect of microtubule drugs on microtubules in quiescent RPE 1 hTert cells. a)** Cell cycle profiles of cycling and quiescent RPE 1 hTert cells treated with DMSO, or microtubule drugs (x-axis). **b and c)** Biochemical partitioning of tubulin into soluble (S) and polymerized (P) in quiescent RPE 1 hTert cells treated with DMSO control, and microtubule drugs at indicated concentrations for 6 (b) and 24h (c). Each soluble tubulin fraction is normalized to the housekeeping gene Glyceraldehyde 3-phosphate dehydrogenase (GAPDH), and each polymerized fraction to the housekeeping gene histone H3 (HISH3). All data are normalized to DMSO-treated control cells. Plotted are average values from 3 biological replicates. **d)** Average number of EB1-positive microtubule plus-tips per cell area in cells treated with microtubule drugs at indicated concentrations (x-axis), and normalized to DMSO-treated control cells (>100 cells per condition). **e)** Representative immunofluorescence images of control and cells treated with indicated microtubule-drugs, for 24 h and stained with anti-EB1 antibody (cyan), anti-β-actin, and Hoechst (red). Bar plots in all panels represent average values, and error bars standard deviations from three independent biological replicates. * p-value<0.05, ** p-value<0.01, *** p-value<0.001 in paired Student T-test compared to DMSO control.

**Supplementary Figure 2. Microtubule damage triggers differential tubulin gene expression across many cancer cell types and *in vivo*. a)** Expression profiles of all detected α- and β-tubulin isoform across a panel of breast, ovarian and endometrial cancer cell lines treated with microtubule poisons for 24h. This dataset is available on GEO database (GEO series GSE50811, GSE50830, and GSE50831[15]). Dendrogram on the left represents Pearson distance between expression profiles. Each column of the heatmap represents differential gene expression in one cell line treated with indicated microtubule drug, marked above the heatmap. Each row represents a gene, labeled on y-axis. Color key is depicted in upper left corner. Data are represented as $Log_2FC$ relative to DMSO control. **b)** Expression profiles of all detected α- and β-tubulin isoforms in heart endothelial cells isolated from control and rats treated with microtubule poisons vinblastine (VIN), vincristine (VCR), or colchicine (COL) at indicated doses (x-axis) for 6 and 24h. This dataset is available on GEO database (GEO series GSE19290). Dendrogram on the left represents Pearson distance between expression profiles. Each column of the heatmap represents differential gene expression in one condition, labeled on the x-axis. Each row represents a gene, labeled on y-axis. Color key is depicted in upper left corner. Data are represented as $Log_2FC$ relative to DMSO control. **c)** Experimental strategy and primer design for RT-qPCR.

**Supplementary Figure 3. Nutrient deprivation regulates the expression of α- and β-tubulin isoforms transcriptionally and post-transcriptionally. a and e)** DGE in control and cells deprived of D-glucose (a, A549 cell line, GEO series GSE56843[26]), or L-glutamine (e, U2OS cell line, GEO series GSE59931). Each column represents DGE in one replicate of control or nutrient deprived cells (x-axis). Each row represents one gene (y-axes). CEM 1 consists of tubulin genes with high Pearson expression correlation. Null module consists of tubulin genes with low Pearson expression correlation. Scales of expression profile Z-scores are depicted above each heatmap. **b and f)** Relative expression of TUBA1A and TUBB unspliced pre-mRNA in control and A549 cells deprived of D-glucose (b), or control and U2OS cells deprived of L-Glutamine for indicated periods of time (f, x-axes). **c and g)** Relative expression of TUBA1A and TUBB mRNA in control and A549 cells deprived of D-glucose (c), or control and U2OS cells deprived of L-Glutamine for indicated periods of time (g, x-axis). All the relative gene expression data are normalized to housekeeping gene GAPDH or RPL19, and to time point 0h (control). **d and h)** Tubulin partitioning to unpolymerized (soluble) and polymerized (pellet), and total tubulin, normalized to loading controls: GAPDH for soluble and total, and HISH3 for pellet, in control and nutrient deprived cells for 24 hours. All data are normalized to DMSO control. Bar plots in all panels represent average values, and error bars standard deviations from three independent biological replicates. * $p<0.05$, ** $p<0.01$, *** $p<0.001$ in Paired Student T-test compared to control treatment.

**Supplementary Figure 4. PI3K inhibitor BKM-120, but not BEZ-235 and GDC-1941, displays off-target effect on microtubules. a)** Quantification of the number of EB-positive microtubule plus-tips per cell area in RPE 1 hTert cells treated with DMSO or indicated concentrations (x-axis) of PI3K inhibitors. Data are normalized to DMSO-treated control cells. Lines represent average number of EB signals per cell area, and shaded area represents standard error of the mean over three independent experiments and >100 cells per condition. **b)** Representative immunofluorescence images of control and cells treated with indicated 1 μM PI3K inhibitors for 6h, and stained with anti-EB1 antibody (cyan), anti-β-actin, and Hoechst (red).

**Supplementary Table 1. RT-qPCR primer sequences.** Listed are target mRNA species, sequences, and orientation of all the primers used for RT-qPCR in this study.

**Supplementary Table 2. Perturbations that trigger tubulin differential expression.** Listed are Rank based on descending Pearson expression-correlation coefficients, GEO identifiers, titles of studies, and observed Pearson expression-correlation coefficients for a subset of tubulin genes across the 417 datasets from human platform in the CLIC-report.

### Supplementary Figure 1

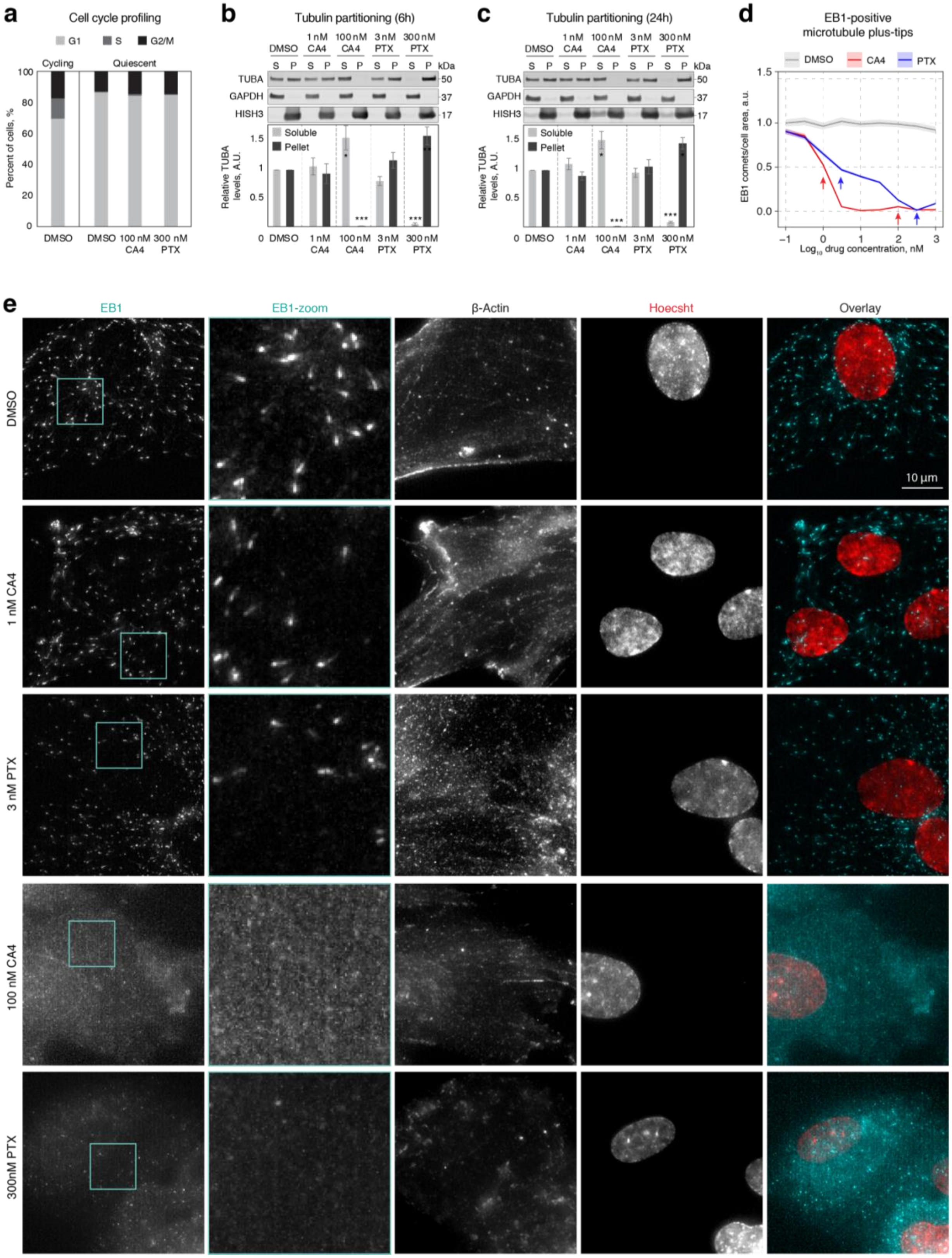

Supplementary Figure 2

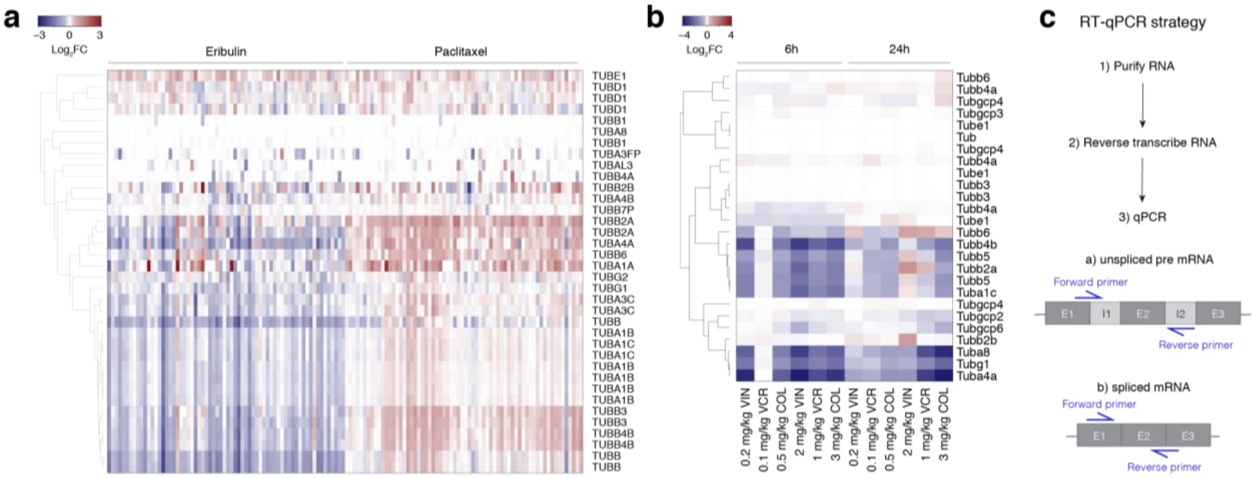

Supplementary Figure 3

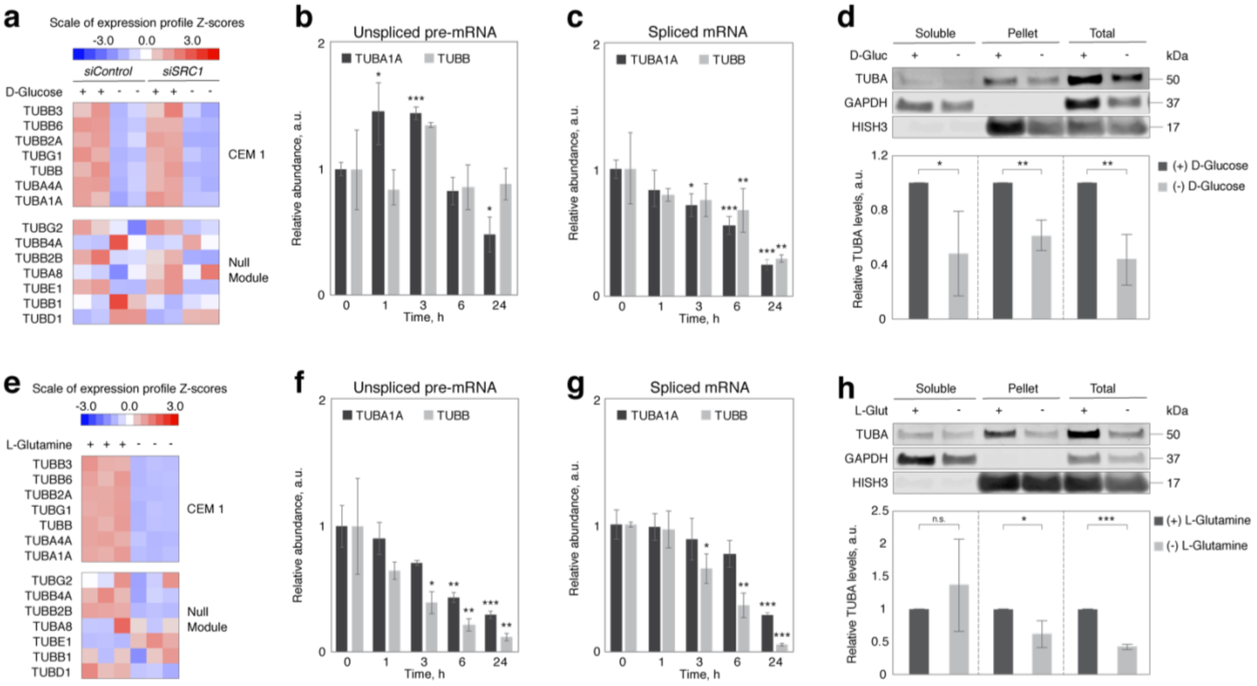

### Supplementary Figure 4

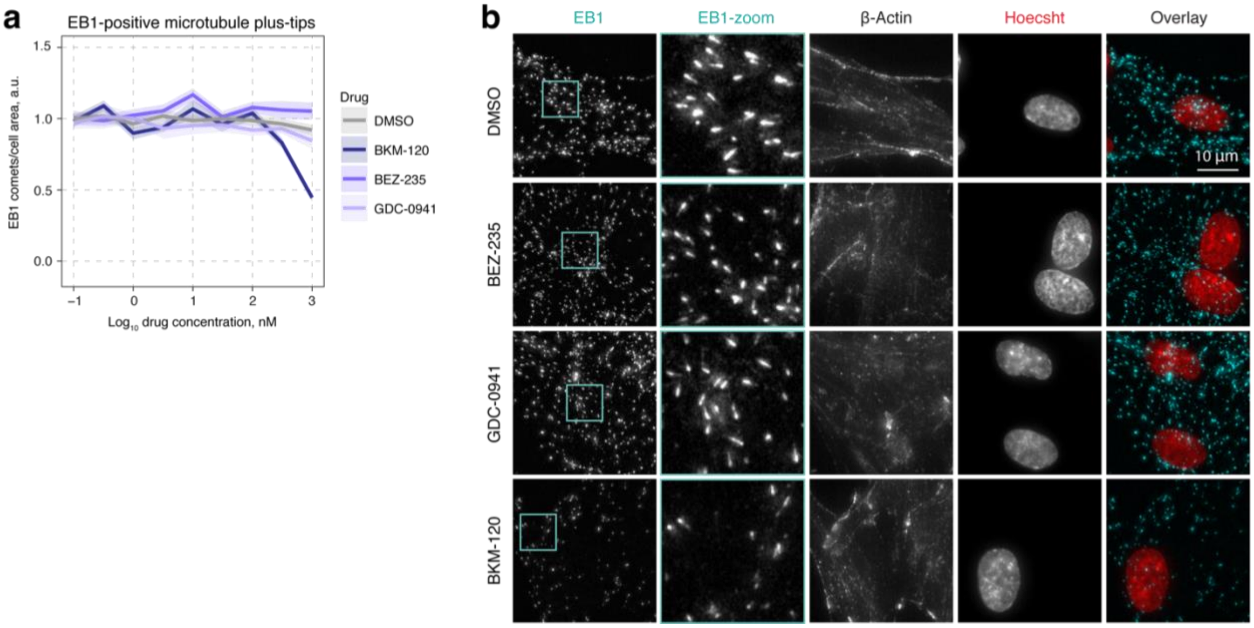

Supplementary Table 1

| **Gene** | **RNA species** | **Orientation** | **Sequence, 5'-3'** |
|---|---|---|---|
| TUBA1A | pre-mRNA | forward | GCAGCATTTGTAGCAGGTGA |
| | | reverse | GCATTGCCAATCTGGACAC |
| | mRNA | forward | CCACAGTCATTGATGAAGTTCG |
| | | reverse | GCTGTGGAAAACCAAGAAGC |
| TUBB | pre-mRNA | forward | CTGGACCGCATCTCTGTGTA |
| | | reverse | GGTTCACGAAAGGGACAAAA |
| | mRNA | forward | GAAGCCACAGGTGGCAAATA |
| | | reverse | CGTACCACATCCAGGACAGA |
| GAPDH | pre-mRNA | forward | GGGAGGTAGAGGGGTGATGT |
| | | reverse | GAGGCAGGGATGATGTTCTG |
| | mRNA | forward | AGCTCATTTCCTGGTATGACA |
| | | reverse | AGGGGAGATTCAGTGTGGTG |
| RPL19 | pre-mRNA | forward | TCCGAGAGGTGAAGGCATAG |
| | | reverse | GCCTCTTCTGAAGCCTGAGC |
| | mRNA | forward | ATCGCCACATGTATCACAGC |
| | | reverse | TTGGTCTCTTCCTCCTTGGAT |

### Supplementary Table 2

| | | | Pearson Correlation Coefficient | | | | | | | | |
|---|---|---|---|---|---|---|---|---|---|---|---|
| **Rank** | **GSE** | **Title of the study** | **TUBB3** | **TUBB6** | **TUBB2A** | **TUBG1** | **TUBB** | **TUBA4A** | **TUBA1A** | **TUBB4B** | **TUBA1C** |
| 1 | GSE59931 | Glutamine deprivation in U2OS cells | 0.98 | 0.99 | 0.99 | 0.99 | 0.99 | 0.98 | 0.99 | 0.99 | 0.91 |
| 2 | GSE36529 | Expression data from CtBP knockdown MCF-7 cells | 0.98 | 0.96 | 0.98 | 0.98 | 0.99 | 0.98 | 0.98 | 0.98 | 0.97 |
| 3 | GSE32158 | Bisphenol A Regulates the Expression of DNA Repair Genes in Human Breast Epithelial Cells (expression data) | 0.98 | 0.98 | 0.98 | 0.96 | 0.97 | 0.98 | 0.95 | 0.97 | 0.97 |
| 4 | GSE23952 | Expression data from TGF-beta treated Panc-1 pancreatic adenocarcinoma cell line | 0.98 | 0.98 | 0.97 | 0.97 | 0.92 | 0.98 | 0.98 | 0.98 | 0.98 |
| 5 | GSE22522 | Comparison of the transcriptome of K-LEC spheroids to control LEC spheroids | 0.98 | 0.96 | 0.96 | 0.98 | 0.97 | 0.92 | 0.97 | 0.98 | 0.96 |
| 6 | GSE56843 | Steroid Receptor Coactivator 1 is an Integrator of Glucose and NAD(+)/NADH Homeostasis | 0.93 | 0.97 | 0.97 | 0.96 | 0.97 | 0.98 | 0.95 | 0.97 | 0.96 |
| 7 | GSE20719 | Gene expression changes upon treatment of T47D breast cancer cells with the Pan-PI3 kinase inhibitor GDC-0941 | 0.95 | 0.96 | 0.97 | 0.98 | 0.94 | 0.96 | 0.93 | 0.98 | 0.93 |
| 8 | GSE4217 | Spheroid Formation and Recovery of Human Foreskin Fibroblasts at Ambient Temperature | 0.97 | 0.96 | 0.96 | 0.92 | 0.90 | 0.94 | 0.94 | 0.97 | 0.96 |
| 9 | GSE58605 | Expression data from A549 cells infected by adenovirus not carrying virus associated sequences in the genome. | 0.94 | 0.92 | 0.94 | 0.94 | 0.93 | 0.95 | 0.96 | 0.96 | 0.57 |
| 10 | GSE46708 | CD24 targets | 0.96 | 0.96 | 0.95 | 0.90 | 0.91 | 0.93 | 0.92 | 0.96 | 0.95 |
| 11 | GSE35428 | Transcriptional profiling of clinically relevant SERMs and SERM/estradiol complexes in a cellular model of breast cancer | 0.95 | 0.95 | 0.92 | 0.95 | 0.92 | 0.93 | 0.91 | 0.96 | 0.95 |
| 12 | GSE7745 | Mapping of HNF4Œ± binding sites, acetylation of histone H3 and expression in Caco2 cells | 0.94 | 0.94 | 0.94 | 0.87 | 0.94 | 0.88 | 0.88 | 0.94 | 0.95 |
| 13 | GSE46924 | 27-Hydroxycholesterol links cholesterol and breast cancer pathophysiology. | 0.90 | 0.93 | 0.92 | 0.95 | 0.93 | 0.90 | 0.84 | 0.93 | 0.88 |
| 14 | GSE16659 | Expression data of HGF/cMET pathway in prostate cancer DU145 cell line | 0.94 | 0.94 | 0.80 | 0.92 | 0.92 | 0.92 | 0.88 | 0.84 | 0.31 |
| 15 | GSE52659 | Expression data from WEEV infected BE(2)-C/m cells | 0.92 | 0.88 | 0.94 | 0.91 | 0.94 | 0.87 | 0.79 | 0.83 | 0.87 |
| 16 | GSE10444 | gene expression levels in long-term cultures of human dental pulp stem cells | 0.83 | 0.88 | 0.90 | 0.92 | 0.93 | 0.89 | 0.86 | 0.94 | 0.81 |
| 17 | GSE4218 | Spheroid Formation and Recovery of Human T98G Glioma Cells at Ambient Temperature | 0.92 | 0.93 | 0.87 | 0.84 | 0.81 | 0.93 | 0.88 | 0.94 | 0.91 |
| 18 | GSE15499 | HDAC5 is a repressor of angiogenesis and determines the angiogenic gene expression pattern of endothelial cells | 0.90 | 0.84 | 0.86 | 0.93 | 0.94 | 0.82 | 0.91 | 0.92 | 0.86 |
| 19 | GSE33143 | Targeted disruption of the BCL9/beta-catenin complex in cancer | 0.91 | 0.87 | 0.89 | 0.89 | 0.87 | 0.90 | 0.81 | 0.83 | 0.16 |
| 20 | GSE42733 | Gene expression profile of Nurse-Like Cells (NLC) derived from chronic lymphocytic leukemia | 0.81 | 0.90 | 0.89 | 0.90 | 0.88 | 0.88 | 0.84 | 0.68 | 0.91 |
| 21 | GSE36085 | Regulation of Autophagy by VEGF-C axis in cancer | 0.91 | 0.87 | 0.90 | 0.90 | 0.86 | 0.92 | 0.72 | 0.91 | 0.78 |
| 22 | GSE43700 | Microarray analysis of IL-10 stimulated adherent peripheral blood mononuclear cells | 0.90 | 0.88 | 0.87 | 0.74 | 0.81 | 0.92 | 0.92 | 0.93 | 0.91 |
| 23 | GSE29625 | Human embryonic stem cells derived from embryos at different stages of development share similar transcription profiles | 0.90 | 0.86 | 0.91 | 0.83 | 0.88 | 0.89 | 0.78 | 0.93 | 0.83 |
| 24 | GSE12098 | Comparison of the migration profile of MSCs | 0.88 | 0.70 | 0.88 | 0.89 | 0.90 | 0.89 | 0.81 | 0.93 | 0.89 |
| 25 | GSE13378 | Exposure of squamous esophageal cell line HET-1A to deoxycholic acid (DCA) | 0.85 | 0.74 | 0.90 | 0.87 | 0.85 | 0.87 | 0.87 | 0.89 | -0.19 |
| 26 | GSE32161 | Microarray analysis of genes associated with cell surface NIS protein levels in breast cancer | 0.88 | 0.81 | 0.91 | 0.90 | 0.79 | 0.79 | 0.87 | 0.88 | 0.87 |
| 27 | GSE16070 | Networking of differentially expressed genes in human MCF7 breast cancer cells resistant to methotrexate | 0.90 | 0.90 | 0.79 | 0.77 | 0.83 | 0.84 | 0.90 | 0.92 | 0.73 |
| 28 | GSE36176 | Gene expression arrays on lung cancer cells exposed to Notch inhibitor | 0.88 | 0.85 | 0.90 | 0.90 | 0.71 | 0.90 | 0.71 | 0.92 | 0.87 |
| 29 | GSE23399 | Gene expression profling of human breast carcinoma-associated fibroblasts treated with paclitaxol or doxorubicin at therapeutically relevant doses | 0.85 | 0.85 | 0.85 | 0.81 | 0.86 | 0.77 | 0.86 | 0.90 | 0.82 |

| | | | | | | | | | | | |
|---|---|---|---|---|---|---|---|---|---|---|---|
| 30 | GSE14001 | PAX2: A Potential Biomarker for Low Malignant Potential Ovarian Tumors and Low-Grade Serous Ovarian Carcinomas | 0.89 | 0.87 | 0.88 | 0.82 | 0.80 | 0.69 | 0.87 | 0.84 | 0.83 |
| 31 | GSE17368 | Epiphyseal cartilage | 0.89 | 0.87 | 0.75 | 0.82 | 0.78 | 0.87 | 0.82 | 0.86 | 0.89 |
| 32 | GSE19495 | Global Gene Expression of Human Hepatoma Cells After Amino Acid Limitation | 0.60 | 0.83 | 0.87 | 0.88 | 0.81 | 0.89 | 0.86 | 0.74 | 0.82 |
| 33 | GSE14773 | Roles of EMT regulator in colon cancer | 0.88 | 0.89 | 0.74 | 0.77 | 0.79 | 0.81 | 0.81 | 0.85 | 0.90 |
| 34 | GSE19136 | Gene expression response to implanted drug (paclitaxel)-eluting or bare metal stents in denuded human LIMA arteries | 0.85 | 0.84 | 0.86 | 0.77 | 0.74 | 0.79 | 0.84 | 0.90 | 0.87 |
| 35 | GSE16356 | Lymphatic endothelial cells (LEC) treated with a MAF-targeted siRNA | 0.84 | 0.78 | 0.81 | 0.75 | 0.85 | 0.77 | 0.87 | 0.84 | 0.48 |
| 36 | GSE9835 | Gene Expression Changes in Response to Baculoviral Vector Transduction of Neuronal Cells In Vitro | 0.62 | 0.87 | 0.82 | 0.79 | 0.89 | 0.87 | 0.76 | 0.87 | 0.68 |
| 37 | GSE17044 | Expression data from androgen treated LNCaP cells | 0.89 | 0.80 | 0.88 | 0.69 | 0.85 | 0.73 | 0.77 | 0.89 | 0.62 |
| 38 | GSE9649 | Expression studies of HMEC exposed to lactic acidosis and hypoxia | 0.86 | 0.74 | 0.85 | 0.78 | 0.81 | 0.86 | 0.72 | 0.87 | 0.80 |
| 39 | GSE31641 | Expression data from treatment of human melanocytes with phenolic compounds | 0.85 | 0.79 | 0.83 | 0.70 | 0.84 | 0.75 | 0.83 | 0.86 | 0.90 |
| 40 | GSE13142 | HepG2/C3A cells cultured for 42 h in complete or leucine-devoid medium | 0.89 | 0.48 | 0.84 | 0.88 | 0.87 | 0.85 | 0.75 | 0.89 | 0.88 |
| 41 | GSE7345 | Germline NRAS mutation causes a novel human autoimmune lymphoproliferative syndrome | 0.86 | 0.66 | 0.73 | 0.83 | 0.86 | 0.83 | 0.75 | 0.88 | 0.87 |
| 42 | GSE34635 | Defining a No Observable Transcriptional Effect Level (NOTEL) for low dose N-OH-PhIP exposures in human BEAS-2B bronchioepithelial cells | 0.84 | 0.82 | 0.82 | 0.79 | 0.50 | 0.86 | 0.86 | 0.86 | 0.81 |
| 43 | GSE5838 | Expression data from transplanted intestine bifor signs of rejection, and when their was signs of rejection | 0.86 | 0.58 | 0.72 | 0.86 | 0.80 | 0.87 | 0.79 | 0.88 | 0.89 |
| 44 | GSE51130 | Using a rhabdomyosarcoma patient-derived xenograft to examine precision medicine approaches and model acquired resistance | 0.88 | 0.57 | 0.85 | 0.76 | 0.80 | 0.84 | 0.79 | 0.89 | 0.76 |
| 45 | GSE34512 | PBEF Knockdown in HMVEC-LBI | 0.87 | 0.88 | 0.84 | 0.84 | 0.84 | 0.80 | 0.39 | 0.89 | 0.46 |
| 46 | GSE42853 | Distinct gene expression profiles associated with the susceptibility of pathogen-specific CD4 T cells to HIV-1 infection | 0.86 | 0.65 | 0.83 | 0.81 | 0.86 | 0.58 | 0.87 | 0.89 | 0.89 |
| 47 | GSE26884 | Bisphenol A Induced the Expression of DNA Repair Genes in Human Breast Epithelial Cells | 0.80 | 0.84 | 0.80 | 0.62 | 0.83 | 0.88 | 0.67 | 0.76 | 0.76 |
| 48 | GSE53731 | Expression data from hepatitis E virus inoculated PLC/PRF/5 cells | 0.77 | 0.85 | 0.75 | 0.85 | 0.73 | 0.65 | 0.83 | 0.78 | 0.82 |
| 49 | GSE8588 | OH-PBDE-induced gene expression profiling in H295R adrenocortical carcinoma cells | 0.87 | 0.52 | 0.85 | 0.69 | 0.83 | 0.82 | 0.85 | 0.88 | 0.82 |
| 50 | GSE12875 | Impaired T-cell function in patients with novel ICOS | 0.86 | 0.52 | 0.87 | 0.83 | 0.82 | 0.77 | 0.75 | 0.88 | 0.83 |
| 51 | GSE45636 | eIF3a in Urinary Bladder Cancer ,Äì in vivo and in vitro insights | 0.87 | 0.84 | 0.81 | 0.54 | 0.86 | 0.81 | 0.68 | 0.83 | 0.72 |
| 52 | GSE49085 | Identification of bone morphogenetic protein (BMP)-7 as a key instructive factor for human epidermal Langerhans cell differentiation and proliferation | 0.84 | 0.82 | 0.84 | 0.86 | 0.76 | 0.60 | 0.68 | 0.85 | 0.81 |
| 53 | GSE31472 | Host cell gene expression in Influenza A/duck/Malaysia/F118/08/2004 (H5N2) infected A549 cells at 2, 4, 6, 8, and 10 hours post infection | 0.80 | 0.84 | 0.84 | 0.56 | 0.86 | 0.80 | 0.69 | 0.88 | -0.37 |
| 54 | GSE9677 | Gene expression profile in HUVECs before and after Angiopoietin stimulation | 0.83 | 0.86 | 0.85 | 0.85 | 0.73 | 0.83 | 0.44 | 0.87 | 0.69 |
| 55 | GSE6400 | Cultured A549 lung cancer cells treated with actinomycin D and sapphyrin PCI-2050 | 0.83 | 0.81 | 0.85 | 0.78 | 0.58 | 0.79 | 0.71 | 0.77 | 0.53 |
| 56 | GSE16089 | Networking of differentially expressed genes in human Saos-2 osteosarcoma cells resistant to methotrexate | 0.85 | 0.73 | 0.36 | 0.85 | 0.86 | 0.86 | 0.85 | 0.89 | 0.89 |
| 57 | GSE15065 | C/EBPbeta-2 regulation of gene expression in MCF10A cells | 0.86 | 0.82 | 0.78 | 0.74 | 0.80 | 0.83 | 0.49 | 0.86 | 0.82 |
| 58 | GSE5110 | 48h Immobilization in human | 0.77 | 0.73 | 0.73 | 0.84 | 0.61 | 0.81 | 0.82 | 0.87 | 0.76 |
| 59 | GSE44540 | Gene expression in hTERT-RPE1 cells with overexpression of MFRP | 0.85 | 0.84 | 0.79 | 0.86 | 0.39 | 0.75 | 0.84 | 0.89 | 0.83 |
| 60 | GSE23764 | Expression data from actomyosin contractility regulated genes | 0.80 | 0.69 | 0.82 | 0.77 | 0.72 | 0.80 | 0.70 | 0.86 | 0.72 |
| 61 | GSE20037 | cdr2 siRNA knockdown during passage through mitosis: HeLa cells, Rat1 wild type and c-myc null cells | 0.80 | 0.46 | 0.82 | 0.79 | 0.79 | 0.82 | 0.80 | 0.70 | 0.47 |

| | | | | | | | | | | | |
|---|---|---|---|---|---|---|---|---|---|---|---|
| 62 | GSE40517 | Selective Requirement for Mediator MED23 in Ras-active Lung Cancer | 0.79 | 0.75 | 0.84 | 0.65 | 0.82 | 0.77 | 0.65 | 0.71 | 0.79 |
| 63 | GSE19510 | Transcriptional response of normal human lung WI-38 fibroblasts to benzo[a]pyrene diol epoxide: a dose-response study | 0.82 | 0.86 | 0.66 | 0.64 | 0.79 | 0.73 | 0.77 | 0.85 | 0.88 |
| 64 | GSE17785 | Endogenous expression of an oncogenic PI3K mutation leads to activated PI3K pathway signaling and an invasive phenotype | 0.83 | 0.82 | 0.81 | 0.51 | 0.78 | 0.81 | 0.65 | 0.86 | 0.80 |
| 65 | GSE29384 | Tetracycline-Inducible Cyr61 effect on LN229 glioma cells | 0.79 | 0.83 | 0.84 | 0.83 | 0.83 | 0.25 | 0.83 | 0.86 | 0.68 |
| 66 | GSE28339 | Gene expression data following Cyclin T2 and Cyclin T1 depletion by shRNA in HeLa cells | 0.83 | 0.81 | 0.53 | 0.54 | 0.83 | 0.83 | 0.76 | 0.83 | 0.65 |
| 67 | GSE2328 | Application of genome-wide expression analysis to human health & disease | 0.77 | 0.78 | 0.76 | 0.36 | 0.83 | 0.80 | 0.81 | 0.85 | 0.78 |
| 68 | GSE33606 | Gene expression changes in human hepatocytes exposed to VX (O-ethyl S-[2-(diisopropylamino)ethyl] methylphosphonothiolate) | 0.82 | 0.77 | 0.81 | 0.53 | 0.84 | 0.75 | 0.54 | 0.86 | 0.84 |
| 69 | GSE37648 | Gene signatures of normal hTERT immortalized ovarian epithelium and fallopian tube epithelium (paired cultures from 2 donor patients) | 0.84 | 0.77 | 0.71 | 0.74 | 0.75 | 0.77 | 0.50 | 0.79 | 0.82 |
| 70 | GSE45417 | Expression data from knockdown of ZXDC1/2 in PMA-treated U937 | 0.79 | 0.74 | 0.75 | 0.79 | 0.74 | 0.81 | 0.45 | 0.80 | 0.85 |
| 71 | GSE37474 | Dexamethasone induced gene expression changes in the human trabecular meshwork | 0.79 | 0.78 | 0.74 | 0.72 | 0.74 | 0.57 | 0.69 | 0.83 | 0.77 |
| 72 | GSE24224 | Analysis of genome-wide methylation and gene expression induced by decitabine treatment in HL60 leukemia cell line | 0.70 | 0.80 | 0.81 | 0.62 | 0.51 | 0.78 | 0.82 | 0.63 | 0.51 |
| 73 | GSE26599 | Gene expression profile in response to doxorubicin-rapamycin combined treatment of HER-2 overexpressing human mammary epithelial cell lines | 0.82 | 0.81 | 0.81 | 0.78 | 0.79 | 0.55 | 0.47 | 0.81 | 0.62 |
| 74 | GSE20125 | Transcriptome analysis of human Wharton‚Äôs jelly stem cells: meta-analysis | 0.75 | 0.73 | 0.70 | 0.80 | 0.77 | 0.48 | 0.72 | 0.84 | 0.81 |
| 75 | GSE47874 | The Heritage (HEalth, RIsk factors, exercise Training And GEnetics) family study | 0.75 | 0.64 | 0.77 | 0.74 | 0.77 | 0.63 | 0.66 | 0.78 | 0.79 |
| 76 | GSE6494 | Expression data from human liver cell line induced by PCB153 | 0.80 | 0.75 | 0.63 | 0.57 | 0.75 | 0.65 | 0.78 | 0.75 | 0.59 |
| 77 | GSE38517 | Expression data from fibroblasts derived from human normal oral mucosa, oral dysplasia and oral squamous cell carcinoma | 0.82 | 0.82 | 0.73 | 0.75 | 0.73 | 0.64 | 0.41 | 0.82 | 0.81 |
| 78 | GSE33243 | Human acute myelogenous leukemia-initiating cells treated with fenretinide | 0.78 | 0.46 | 0.66 | 0.73 | 0.68 | 0.76 | 0.81 | 0.80 | 0.73 |
| 79 | GSE22533 | Breast cancer cells resistant to hormone deprivation maintain an estrogen receptor alpha-dependent, E2F-directed transcriptional program | 0.81 | 0.83 | 0.78 | 0.82 | 0.81 | 0.41 | 0.40 | 0.85 | 0.85 |
| 80 | GSE30494 | Expression data in cancer cell lines using Affymetrix GeneChip | 0.63 | 0.79 | 0.80 | 0.79 | 0.36 | 0.69 | 0.76 | 0.79 | -0.37 |
| 81 | GSE4824 | Analysis of lung cancer cell lines | 0.77 | 0.80 | 0.69 | 0.45 | 0.80 | 0.60 | 0.70 | 0.78 | 0.81 |
| 82 | GSE15372 | Expression data from A2780 (cisplatin-sensitive) and Round5 A2780 (cisplatin-resistant) cell lines. | 0.53 | 0.78 | 0.82 | 0.76 | 0.79 | 0.48 | 0.64 | 0.81 | 0.47 |
| 83 | GSE16524 | Expression data from skin fibroblasts derived from Setleis Syndrome patients and normal controls | 0.81 | 0.68 | 0.70 | 0.73 | 0.72 | 0.66 | 0.50 | 0.84 | 0.72 |
| 84 | GSE13054 | Genes upregulated by HLX | 0.82 | 0.37 | 0.70 | 0.69 | 0.78 | 0.75 | 0.68 | 0.80 | 0.80 |
| 85 | GSE12274 | Mesenchymal Stromal Cells of Different Donor Age | 0.80 | 0.82 | 0.53 | 0.71 | 0.77 | 0.60 | 0.56 | 0.84 | 0.79 |
| 86 | GSE35170 | Expression data from U87-2M1 glioma cells transduced with baculoviral control decoy vector or baculoviral miR-10b decoy vector | 0.78 | 0.71 | 0.68 | 0.38 | 0.72 | 0.73 | 0.77 | 0.80 | 0.41 |
| 87 | GSE20540 | Gene expression profiles of myeloma cells interacting with bone marrow stromal cells in vitro | 0.72 | 0.68 | 0.73 | 0.75 | 0.76 | 0.34 | 0.76 | 0.36 | 0.37 |
| 88 | GSE4289 | Host transcriptome changes associated with episome loss and selection of keratinocytes containing integrated HPV16 | 0.77 | 0.80 | 0.80 | 0.62 | 0.57 | 0.78 | 0.39 | 0.71 | 0.79 |
| 89 | GSE11352 | Timecourse of estradiol (10nM) exposure in MCF7 breast cancer cells. | 0.73 | 0.66 | 0.77 | 0.69 | 0.61 | 0.58 | 0.67 | 0.77 | 0.56 |
| 90 | GSE39999 | Filarial nematode AsnRS interacts with interleukin 8 receptors in iDCs but causes different gene expression patterns compared to iDCs stimulated by interleukin 8. | 0.72 | 0.75 | 0.74 | 0.72 | 0.73 | 0.58 | 0.66 | 0.80 | 0.81 |

| | | | | | | | | | | | |
|---|---|---|---|---|---|---|---|---|---|---|---|
| 91 | GSE11208 | Chronic nicotine exposure (kuo-affy-human-232930) | 0.70 | 0.72 | 0.47 | 0.75 | 0.72 | 0.67 | 0.65 | 0.68 | 0.70 |
| 92 | GSE14986 | Antiestrogen-resistant subclones of MCF-7 human breast cancer cells are derived from a common clonal drug-resistant progenitor | 0.70 | 0.57 | 0.66 | 0.62 | 0.74 | 0.73 | 0.67 | 0.74 | 0.62 |
| 93 | GSE16538 | Genome-wide gene expression profile analysis in pulmonary sarcoidosis | 0.75 | 0.67 | 0.71 | 0.70 | 0.73 | 0.68 | 0.42 | 0.70 | 0.71 |
| 94 | GSE18182 | Expression profile of lung adenocarcinoma, A549 cells following targeted depletion of non metastatic 2 (NME2/NM23 H2) | 0.74 | 0.62 | 0.46 | 0.70 | 0.81 | 0.71 | 0.61 | 0.75 | -0.05 |
| 95 | GSE32892 | A genome-wide and dose-dependent inhibition map of androgen receptor binding by small molecules reveals its regulatory program upon antagonism | 0.74 | 0.78 | 0.76 | 0.81 | 0.76 | 0.68 | 0.11 | 0.82 | 0.78 |
| 96 | GSE40220 | INTESTINAL FILTER FOR USE IN OESOPHAGEAL CANCER RESEARCH | 0.73 | 0.80 | 0.74 | 0.75 | 0.67 | 0.80 | 0.14 | 0.74 | 0.75 |
| 97 | GSE33146 | Expression data from DKAT breast cancer cell line pre- and post-EMT | 0.78 | 0.72 | 0.74 | 0.74 | 0.52 | 0.69 | 0.45 | 0.63 | -0.09 |
| 98 | GSE17624 | Expression data from human Ishikawa cells treated with Bisphenol A | 0.73 | 0.78 | 0.60 | 0.61 | 0.75 | 0.72 | 0.43 | 0.80 | 0.39 |
| 99 | GSE17090 | Expression data from human adipose stem cells expanded in allogeneic human serum and fetal bovine serum | 0.77 | 0.72 | 0.75 | 0.72 | 0.50 | 0.59 | 0.52 | 0.78 | 0.80 |
| 100 | GSE16054 | Transient expression of misfolded surfactant protein C | 0.56 | 0.56 | 0.79 | 0.76 | 0.75 | 0.70 | 0.41 | 0.76 | 0.69 |
| 101 | GSE16715 | Expression profiling in Williams-Beuren Syndrome patient fibroblast cell lines | 0.76 | 0.61 | 0.62 | 0.52 | 0.73 | 0.58 | 0.70 | 0.78 | 0.73 |
| 102 | GSE9593 | Cellular Aging of Mesenchymal Stem Cells | 0.75 | 0.77 | 0.40 | 0.63 | 0.78 | 0.74 | 0.43 | 0.79 | 0.73 |
| 103 | GSE7846 | Differentially expressed genes in HEECs of eutopic endometrium of patients with endometriosis compared with control | 0.72 | 0.69 | 0.77 | 0.62 | 0.64 | 0.35 | 0.67 | 0.80 | 0.70 |
| 104 | GSE13671 | Expression data from mammary epithelial cells from BRCA1 mutation carriers and non BRCA1 mutation carriers | 0.75 | 0.70 | 0.54 | 0.61 | 0.51 | 0.72 | 0.65 | 0.67 | 0.75 |
| 105 | GSE8045 | Gene Expression Profiling in A549 Lung Cancer Cell Line Following siRNA Mediated Knock-down of ALDH1A1 and ALDH3A1 | 0.78 | 0.75 | 0.70 | 0.72 | 0.67 | 0.16 | 0.70 | 0.80 | 0.81 |
| 106 | GSE11550 | Hs 294T Cells Treated with Elesclomol Alone or in Combination with Paclitaxel Compared to DMSO Treated | 0.70 | 0.48 | 0.76 | 0.49 | 0.74 | 0.65 | 0.66 | 0.81 | 0.54 |
| 107 | GSE6869 | Expression data from human liver cell line induced by PCB77 | 0.72 | 0.76 | 0.51 | 0.66 | 0.70 | 0.66 | 0.45 | 0.75 | 0.64 |
| 108 | GSE51549 | All trans-retinoic acid (ATRA) re-differentiate early transformed breast epithelial cells to normal. | 0.79 | 0.69 | 0.73 | 0.40 | 0.46 | 0.70 | 0.67 | 0.80 | 0.66 |
| 109 | GSE32939 | CD4 on human monocytes | 0.80 | 0.42 | 0.59 | 0.69 | 0.41 | 0.78 | 0.77 | 0.83 | 0.73 |
| 110 | GSE16547 | KSHV Manipulates Notch Signaling by Upregulating Dll4 and JAG1 to Alter Cell Cycle Gene Expression in LECs | 0.75 | 0.70 | 0.74 | 0.71 | 0.73 | 0.75 | 0.06 | 0.77 | 0.75 |
| 111 | GSE14807 | Investigation of over-expressing Annexin receptor cell line with and without agonists | 0.57 | 0.75 | 0.69 | 0.75 | 0.60 | 0.66 | 0.76 | 0.59 | 0.37 |
| 112 | GSE18161 | Washing scaling of microarray expression | 0.63 | 0.74 | 0.66 | 0.42 | 0.71 | 0.53 | 0.69 | 0.72 | 0.71 |
| 113 | GSE9361 | Functional interaction between a PIP2 novel polyA polymerase and type 1 PIPKIalpha | 0.71 | 0.73 | 0.62 | 0.73 | 0.08 | 0.75 | 0.75 | 0.78 | 0.71 |
| 114 | GSE33455 | Expression data from docetaxel-resistant prostate cancer cell lines | 0.52 | 0.77 | 0.68 | 0.76 | 0.78 | 0.66 | 0.18 | 0.70 | -0.16 |
| 115 | GSE32984 | Gene expression profiling of Human Umbilical Vein Endothelial Cells (HUVEC) after treatment with Erg or control antisense (GeneBloc) | 0.66 | 0.72 | 0.63 | 0.47 | 0.75 | 0.70 | 0.42 | 0.75 | 0.65 |
| 116 | GSE30038 | Programming human pluripotent stem cells into adipocytes [Affymetrix] | 0.68 | 0.75 | 0.74 | 0.39 | 0.75 | 0.35 | 0.74 | 0.69 | 0.62 |
| 117 | GSE35957 | Effects of Cellular Senescence on Human Mesenchymal Stem Cells | 0.63 | 0.62 | 0.64 | 0.73 | 0.70 | 0.50 | 0.49 | 0.77 | 0.76 |
| 118 | GSE10934 | Human sclera | 0.71 | 0.74 | 0.69 | 0.75 | 0.73 | 0.02 | 0.68 | 0.79 | 0.76 |
| 119 | GSE18892 | Silencing of AEBP1 in U87MG glial cells and Chip-chIP with AEBP1 antibody | 0.78 | 0.71 | 0.70 | 0.58 | 0.45 | 0.65 | 0.57 | 0.78 | 0.75 |
| 120 | GSE47873 | Gene expression profiles MCF-10A cells overexpressing MBD2 | 0.79 | 0.76 | 0.39 | 0.74 | 0.69 | 0.75 | 0.17 | 0.78 | 0.80 |
| 121 | GSE31469 | Host cell gene expression in Influenza A/WSN/33 (H1N1) infected A549 cells at 2, 4 and 6 hours post infection | 0.71 | 0.32 | 0.64 | 0.52 | 0.67 | 0.72 | 0.71 | 0.77 | 0.25 |
| 122 | GSE28448 | Antagonistic regulation of EMT by TIF1Œ≥ and Smad4 in mammary epithelial cells | 0.56 | 0.74 | 0.73 | 0.59 | 0.65 | 0.73 | 0.30 | 0.66 | 0.71 |

| | | | | | | | | | | | |
|---|---|---|---|---|---|---|---|---|---|---|---|
| 123 | GSE18866 | Expression data from doxycylin-inducible miR-15a/16-1 and empty vector (EV) expression in a 13q14-\- cell line | 0.68 | 0.71 | 0.44 | 0.72 | 0.30 | 0.74 | 0.70 | 0.76 | 0.28 |
| 124 | GSE28829 | Gene Expression in early and advanced atherosclerotic plaque from human carotid | 0.71 | 0.72 | 0.68 | 0.66 | 0.45 | 0.44 | 0.59 | 0.76 | 0.65 |
| 125 | GSE9768 | Identification of genes modulated by acid and bile in a Barrett's oesophagus cell line | 0.75 | 0.69 | 0.11 | 0.74 | 0.75 | 0.71 | 0.52 | 0.76 | 0.47 |
| 126 | GSE7700 | Genes regulated by YAP in normal breast cell line and breast cancer cell lines | 0.76 | 0.74 | 0.74 | 0.76 | 0.73 | -0.24 | 0.76 | 0.79 | 0.80 |
| 127 | GSE16962 | Effect of mir-210 overexpression or down-modulation on human umbilical vein cells | 0.72 | 0.70 | 0.75 | 0.72 | 0.31 | 0.75 | 0.31 | 0.73 | 0.72 |
| 128 | GSE21570 | Frizzled 4 regulates stemness and invasiveness of migrating glioma cells established by serial in vivo intracranial transplantation | 0.66 | 0.77 | 0.75 | 0.63 | 0.67 | 0.72 | 0.36 | 0.65 | 0.71 |
| 129 | GSE25746 | Integrated, genome-wide screening for hypomethylated oncogenes in salivary gland adenoid cystic carcinoma | 0.72 | 0.73 | 0.12 | 0.69 | 0.63 | 0.57 | 0.76 | 0.75 | 0.76 |
| 130 | GSE27200 | Expression data from Sotos syndrome patients and controls | 0.73 | 0.66 | 0.54 | 0.58 | 0.63 | 0.53 | 0.54 | 0.74 | 0.78 |
| 131 | GSE61352 | Expression data from normal urothelial cells with exogenous expression of mutant FGFR3 | 0.32 | 0.75 | 0.78 | 0.70 | 0.63 | 0.78 | 0.61 | 0.48 | 0.72 |
| 132 | GSE7824 | Zonal Heterogeneity for Gene Expression in Human Pancreatic Carcinoma Growing in the Pancreas of Nude Mice | 0.69 | 0.68 | 0.62 | 0.62 | 0.54 | 0.61 | 0.45 | 0.68 | 0.62 |
| 133 | GSE17466 | Gene expression in iPrEC cell line transfected with wild-type USP2A, mutant USP2A, or empty vector. | 0.73 | 0.62 | 0.39 | 0.68 | 0.67 | 0.69 | 0.43 | 0.76 | 0.67 |
| 134 | GSE29265 | Sporadic vs. Post-Chernobyl Papillary vs. Anaplastic Thyroid Cancers | 0.65 | 0.66 | 0.64 | 0.57 | 0.64 | 0.44 | 0.59 | 0.48 | 0.74 |
| 135 | GSE42762 | FOXO3a Is A Major Target Of Inactivation By PI3K/AKT Signaling In Aggressive Neuroblastoma | 0.71 | 0.68 | 0.67 | 0.75 | 0.72 | 0.57 | 0.09 | 0.79 | 0.53 |
| 136 | GSE30336 | Expression analysis of 52 glioma clinical samples (36 CIMP+ and 16 CIMP-) and 6 cell line samples | 0.68 | 0.56 | 0.79 | 0.76 | 0.63 | 0.40 | 0.62 | 0.66 | 0.42 |
| 137 | GSE33168 | The amniotic fluid transcriptome: a source of novel information about human fetal development | 0.48 | 0.66 | 0.72 | 0.53 | 0.65 | 0.47 | 0.67 | 0.71 | 0.18 |
| 138 | GSE6460 | Human mesenchymal stem cells | 0.67 | 0.67 | 0.74 | 0.65 | 0.73 | 0.59 | 0.12 | 0.68 | 0.49 |
| 139 | GSE57441 | Genes expression of cervical squamous cell carcinoma, CaSki cells, treated with or without recombinant TGF-b1 (2 ng/mL) | 0.66 | 0.73 | 0.70 | -0.02 | 0.71 | 0.67 | 0.70 | 0.23 | -0.28 |
| 140 | GSE40611 | Gene expression data of parotid tissue from Primary Sjogren‚Äôs Syndrome and controls | 0.67 | 0.55 | 0.56 | 0.58 | 0.61 | 0.70 | 0.47 | 0.74 | 0.73 |
| 141 | GSE6679 | Staufen1 regulates a variety of mammalian transcripts | 0.68 | 0.76 | 0.60 | 0.73 | 0.30 | 0.77 | 0.53 | 0.38 | -0.11 |
| 142 | GSE36970 | KDM4B- and KDM6B-regulated genes in human mesenchymal stem cell osteogenic differentiation | 0.71 | 0.67 | 0.74 | 0.28 | 0.49 | 0.53 | 0.67 | 0.79 | 0.79 |
| 143 | GSE30391 | Expression data from human Wharton's jelly stem cells | 0.65 | 0.69 | 0.64 | 0.47 | 0.70 | 0.43 | 0.48 | 0.73 | 0.66 |
| 144 | GSE7807 | Leukotriene D4 induces gene expression in human monocytes through cysteinyl leukotriene type I receptor. | 0.69 | 0.64 | 0.74 | 0.43 | 0.71 | 0.34 | 0.59 | 0.65 | 0.57 |
| 145 | GSE35382 | Comparison of gene expression profiles of HT29 cells treated with Instant Caffeinated Coffee or Caffeic Acid versus control. | 0.62 | 0.54 | 0.52 | 0.73 | 0.65 | 0.68 | 0.46 | 0.65 | 0.55 |
| 146 | GSE48616 | Expression data from hBMSCs cultured on PLLA nanofibers | 0.73 | 0.74 | 0.51 | 0.73 | 0.65 | -0.10 | 0.72 | 0.78 | 0.77 |
| 147 | GSE8597 | Gene expression analysis of hormone treated MCF7 breast cancer cells in the presence or absence of cycloheximide | 0.72 | 0.70 | -0.29 | 0.73 | 0.72 | 0.72 | 0.66 | 0.76 | 0.76 |
| 148 | GSE47920 | Expression data from T lymphocytes derived from T-iPS and peripheral blood | 0.70 | 0.66 | 0.25 | 0.70 | 0.74 | 0.33 | 0.58 | 0.73 | 0.67 |
| 149 | GSE9212 | Overexpression of lung developmental transcription factors TTF-1, NKX2-8 and PAX9 | 0.70 | 0.69 | 0.18 | 0.53 | 0.61 | 0.63 | 0.58 | 0.69 | 0.48 |
| 150 | GSE19123 | Lactic acidosis triggers starvation response with distinct metabolic profiles | 0.66 | 0.63 | 0.53 | 0.63 | 0.62 | 0.36 | 0.48 | 0.67 | 0.68 |
| 151 | GSE25148 | Changes in gene expression in HEK-TLR2 cells in response to Helicobacter pylori lipopolysaccharide | 0.68 | 0.66 | 0.59 | 0.59 | 0.55 | 0.52 | 0.44 | 0.70 | 0.61 |
| 152 | GSE13330 | Senescent Stromal-Derived Osteopontin Promotes Preneoplastic Cell Growth | 0.59 | 0.61 | 0.70 | 0.57 | 0.04 | 0.70 | 0.68 | 0.75 | 0.75 |
| 153 | GSE57634 | Molecular effects of EtOH and Nicotine on normal human oral keratinocytes | 0.61 | 0.28 | 0.68 | 0.60 | 0.58 | 0.53 | 0.60 | 0.66 | 0.23 |

| | | | | | | | | | | | |
|---|---|---|---|---|---|---|---|---|---|---|---|
| 154 | GSE26704 | p38alfa and ATF2 act differentially depending on DUSP1 expression in NCSCL in response to cisplatin | 0.70 | 0.66 | 0.52 | 0.51 | 0.54 | 0.61 | 0.45 | 0.73 | 0.68 |
| 155 | GSE37136 | Gene expression profiling of enforced HOXA1 expression in melanoma cell line | 0.73 | 0.72 | 0.73 | 0.74 | 0.74 | 0.74 | -0.55 | 0.77 | 0.69 |
| 156 | GSE27316 | Effects of long dsRNA expression in HeLa and HEK293 cells | 0.73 | 0.72 | 0.32 | 0.71 | 0.71 | 0.73 | -0.09 | 0.77 | 0.69 |
| 157 | GSE60771 | Testing gene expression changes in VCaP upon depletion of the mutated ETS transcription factor ERG | 0.66 | 0.50 | 0.62 | 0.67 | 0.71 | 0.67 | -0.02 | 0.71 | 0.67 |
| 158 | GSE11418 | Passage dependent gene expression in normal human dermal fibroblasts | 0.66 | 0.61 | 0.28 | 0.54 | 0.63 | 0.43 | 0.68 | 0.75 | 0.67 |
| 159 | GSE31534 | Gene expression profile in A375 melanoma cells after 45 functionally important molecules were knocked down using siRNA | 0.67 | 0.67 | 0.58 | 0.66 | 0.67 | 0.56 | -0.04 | 0.71 | 0.67 |
| 160 | GSE7268 | Cryptosporidium infection of human intestinal tissues causes increased expression of Osteoprotegerin | 0.66 | 0.61 | 0.49 | 0.61 | 0.45 | 0.60 | 0.47 | 0.59 | 0.55 |
| 161 | GSE14334 | Transcriptomic analysis of human lung development | 0.60 | 0.65 | 0.63 | 0.61 | 0.65 | -0.09 | 0.65 | 0.68 | 0.71 |
| 162 | GSE13491 | Therapeutic efficacy of human umbilical cord blood-derived mesenchymal stem cells in myocardial repair after infarction | 0.71 | 0.63 | 0.66 | 0.66 | 0.69 | 0.63 | -0.29 | 0.75 | 0.72 |
| 163 | GSE42924 | Expression data from immature dendritic cells (iDC) expressing HIV-1 Tat alleles and mutants | 0.58 | 0.67 | 0.49 | 0.28 | 0.61 | 0.50 | 0.55 | 0.71 | 0.61 |
| 164 | GSE12198 | Primary NKcells vs. NKAES-derived NK cells vs. NKcells stimulated by low/high dose IL2 after 7days of culture | 0.67 | 0.29 | 0.33 | 0.68 | 0.67 | 0.50 | 0.54 | 0.74 | 0.73 |
| 165 | GSE20538 | Gene expression profiles of fibroblasts from MCT8 patients | 0.69 | 0.63 | 0.36 | 0.68 | 0.62 | 0.43 | 0.25 | 0.73 | 0.71 |
| 166 | GSE47746 | Gene expression data of fibroblasts transduced with LacZ or p63+KLF4. | 0.71 | 0.71 | 0.53 | 0.68 | 0.64 | -0.06 | 0.45 | 0.76 | 0.64 |
| 167 | GSE23586 | Altered gene expressions of leukocyte transendothelial migration and cell communication pathways in periodontitis-affected gingival tissues | 0.67 | 0.31 | 0.69 | 0.70 | 0.52 | 0.72 | 0.45 | 0.79 | 0.64 |
| 168 | GSE11407 | SCFA-hexosamine scaffold | 0.50 | 0.61 | 0.66 | 0.44 | 0.50 | 0.46 | 0.48 | 0.72 | 0.65 |
| 169 | GSE4984 | Monocyte Derived Dendritic Cell Maturation | 0.59 | 0.56 | 0.17 | 0.65 | 0.64 | 0.49 | 0.59 | 0.52 | 0.38 |
| 170 | GSE45426 | Transcriptional events in human skeletal muscle at the outset of concentric resistance exercise training | 0.66 | 0.57 | 0.65 | 0.36 | 0.62 | 0.19 | 0.59 | 0.68 | 0.73 |
| 171 | GSE20433 | Effects of T350 phospho- on gene repression activity of EZH2 | 0.70 | 0.59 | 0.65 | 0.66 | -0.23 | 0.68 | 0.58 | 0.76 | 0.58 |
| 172 | GSE39762 | Genome Wide Profiling of p53 Response to Differentiation or DNA Damage of Human Embryonic Stem Cells | 0.71 | 0.66 | 0.59 | -0.22 | 0.53 | 0.65 | 0.70 | 0.73 | 0.65 |
| 173 | GSE55609 | Human meningioma culture | 0.69 | 0.55 | 0.63 | 0.56 | 0.39 | 0.56 | 0.24 | 0.70 | 0.70 |
| 174 | GSE7888 | Expression data from human mesenchymal stem cells (six batches) | 0.63 | 0.60 | 0.42 | 0.50 | 0.49 | 0.52 | 0.46 | 0.67 | 0.64 |
| 175 | GSE34112 | Effect of NO-sulindac treatment on hypoxic prostate cancer cells | 0.71 | 0.62 | 0.40 | -0.04 | 0.57 | 0.70 | 0.61 | 0.71 | 0.63 |
| 176 | GSE11869 | The genomic response of a human uterine endometrial adenocarcinoma cell line to 17alpha-ethynyl estradiol. | 0.51 | 0.60 | 0.59 | 0.42 | 0.34 | 0.60 | 0.46 | 0.45 | 0.12 |
| 177 | GSE7224 | Gene expression in tonsil and oral epithelia | 0.65 | 0.36 | 0.69 | 0.64 | -0.05 | 0.68 | 0.56 | 0.67 | 0.71 |
| 178 | GSE53059 | Human Subperitoneal Fibroblasts and Cancer Cell Interaction Creates Microenvironment Enhancing Tumor Progression and Metastasis | 0.66 | 0.65 | 0.68 | 0.65 | 0.44 | -0.20 | 0.62 | 0.72 | 0.70 |
| 179 | GSE27041 | OXPHOS complex I deficiency leads to transcriptional changes of the Nrf2-Keap1 pathway and selenoproteins. | 0.58 | 0.59 | 0.29 | 0.63 | 0.55 | 0.34 | 0.49 | 0.64 | 0.51 |
| 180 | GSE4587 | Whole-genome expression profiling of melanoma progression. | 0.69 | 0.67 | 0.39 | 0.66 | 0.65 | 0.51 | -0.09 | 0.72 | 0.69 |
| 181 | GSE11622 | Molecular Analysis of the Vaginal Response to Estrogens in the Ovariectomized Rat and Postmenopausal Woman | 0.68 | 0.61 | 0.45 | 0.34 | 0.22 | 0.66 | 0.51 | 0.73 | 0.66 |
| 182 | GSE42253 | Gene expression data from T cells and NK cells with and without treatment with Hsp90 inhibitor (Geldanamycin) | 0.71 | 0.70 | 0.61 | -0.02 | 0.63 | 0.68 | 0.19 | 0.75 | 0.72 |
| 183 | GSE4237 | Hussaini-2R01NS035122-06A1 | 0.60 | 0.57 | 0.61 | 0.63 | 0.60 | 0.54 | -0.11 | 0.72 | 0.43 |
| 184 | GSE24235 | Skeletal muscle gene expression in response to resistance exercise: sex specific regulation | 0.56 | 0.46 | 0.55 | 0.41 | 0.65 | 0.36 | 0.42 | 0.56 | 0.53 |
| 185 | GSE30806 | Expression profiling in VCP-associated myopathy | 0.59 | 0.52 | 0.50 | 0.25 | 0.48 | 0.47 | 0.64 | 0.55 | 0.61 |

| | | | | | | | | | | |
|---|---|---|---|---|---|---|---|---|---|---|
| 186 | GSE15543 | Meta analysis of gene expression in human islets after in vitro expansion. | 0.68 | 0.50 | 0.57 | 0.55 | 0.52 | 0.16 | 0.39 | 0.70 | 0.64 |
| 187 | GSE33643 | Comparison of gene expression alterations induced by distinct PI3K inhibitors | 0.46 | 0.54 | 0.66 | 0.62 | 0.33 | 0.33 | 0.41 | 0.64 | 0.61 |
| 188 | GSE46914 | IDENTIFICATION OF BIOMARKERS OF RESPONSE TO IFNG DURING ENDOTOXIN TOLERANCE: APPLICATION TO SEPTIC SHOCK | 0.53 | 0.57 | 0.64 | 0.48 | 0.42 | 0.09 | 0.62 | 0.52 | 0.54 |
| 189 | GSE36895 | Molecular Genetic Classification of clear-cell Renal Cell Carcinoma (ccRCC) based on the Gene Expression Profiling of Tumors and Tumorgrafts deficient for BAP1 or PBRM1 | 0.65 | 0.55 | 0.41 | 0.46 | 0.50 | 0.47 | 0.41 | 0.58 | 0.59 |
| 190 | GSE44596 | The effect of Sparstolonin B (SsnB) on gene expression in HCAECs | 0.67 | 0.65 | 0.63 | 0.65 | 0.65 | 0.68 | -0.61 | 0.71 | 0.68 |
| 191 | GSE57463 | SOX9 overexpression in melanoma | 0.66 | 0.68 | -0.54 | 0.68 | 0.59 | 0.62 | 0.61 | 0.72 | 0.67 |
| 192 | GSE19315 | Global transcriptional response of macrophage-like THP-1 cells | 0.65 | 0.66 | -0.36 | 0.69 | 0.57 | 0.63 | 0.51 | 0.69 | 0.73 |
| 193 | GSE16354 | Infection of Lymphatic and Blood Vessel Endothelial Cells (LEC and BEC) with KSHV | 0.64 | 0.47 | 0.33 | 0.56 | 0.57 | 0.36 | 0.38 | 0.64 | 0.69 |
| 194 | GSE13487 | Antitumor efficacy of RAF inhibitor GDC-0879 involving BRAFV600E mutational status and ERK/MAPK pathway suppression | 0.61 | 0.42 | 0.59 | 0.45 | 0.58 | 0.55 | 0.09 | 0.58 | 0.18 |
| 195 | GSE23994 | A molecular signature of normal breast epithelial and stromal cells from Li-Fraumeni syndrome mutation carriers | 0.66 | 0.65 | 0.62 | 0.57 | 0.65 | -0.50 | 0.63 | 0.73 | 0.70 |
| 196 | GSE36807 | Genome-wide analysis of Crohn's disease and ulcerative colitis biopsy samples. | 0.36 | 0.52 | 0.43 | 0.56 | 0.49 | 0.64 | 0.42 | 0.23 | 0.59 |
| 197 | GSE23849 | The ERAD inhibitor Eeyarestatin I is a bifunctional compound with an ER localizing domain and a p97/VCP inhibitory group | 0.68 | 0.68 | 0.39 | -0.08 | 0.50 | 0.49 | 0.65 | 0.72 | 0.63 |
| 198 | GSE33301 | Expression data from control and COUP-TFII siRNA treated HUVEC cells | 0.74 | 0.51 | 0.73 | 0.52 | 0.62 | 0.69 | 0.12 | 0.77 | 0.69 |
| 199 | GSE14897 | Highly Efficient generation of Human Hepatic Cells from Induced Pluripotent Stem Cells. | 0.63 | -0.08 | 0.63 | 0.59 | 0.58 | 0.62 | 0.27 | 0.70 | 0.66 |
| 200 | GSE43489 | Expression profile from PCE-Epi and derived cell lines | 0.82 | 0.67 | 0.79 | 0.81 | 0.80 | 0.60 | 0.49 | 0.75 | 0.79 |
| 201 | GSE46268 | Gene expression profile of human monocytes stimulated with all-trans retinoic acid (ATRA) or 1,25a-dihydroxyvitamin D3 (1,25D3) | 0.61 | 0.51 | -0.21 | 0.60 | 0.59 | 0.63 | 0.47 | 0.66 | 0.57 |
| 202 | GSE13376 | Exposure of Barrett's associated adenocarcinoma cell lines SKGT4 to deoxycholic acid (DCA) | 0.64 | 0.57 | -0.49 | 0.65 | 0.63 | 0.62 | 0.58 | 0.61 | -0.15 |
| 203 | GSE27850 | HAV acute infection in chimpanzees | 0.60 | 0.35 | 0.52 | 0.27 | 0.51 | 0.43 | 0.52 | 0.60 | 0.60 |
| 204 | GSE59412 | A Systems Biology Approach identified different regulatory networks targeted by KSHV miR-K12-11 in B cells and endothelial cells | 0.64 | 0.56 | 0.57 | 0.66 | 0.64 | -0.45 | 0.57 | 0.67 | 0.64 |
| 205 | GSE3397 | RSV gene expression | 0.65 | 0.63 | 0.39 | 0.14 | 0.16 | 0.65 | 0.57 | 0.70 | 0.22 |
| 206 | GSE52674 | expression data of miR-21 knockdown in MCF-7 | 0.51 | 0.66 | 0.65 | 0.65 | 0.58 | 0.63 | -0.51 | 0.59 | 0.64 |
| 207 | GSE49967 | Variability in functional p53 reactivation by Prima-1Met/APR-246 in Ewing sarcoma | 0.64 | 0.58 | 0.67 | 0.44 | 0.72 | 0.48 | 0.49 | 0.58 | 0.69 |
| 208 | GSE55859 | Gene expression profile of TRAIL-sensitive and -resistant H460 cells | 0.67 | 0.66 | 0.66 | 0.67 | 0.67 | 0.63 | -0.84 | 0.71 | -0.18 |
| 209 | GSE13637 | Influenza virus infected HUVEC | 0.39 | 0.59 | 0.40 | 0.15 | 0.58 | 0.55 | 0.46 | 0.60 | 0.37 |
| 210 | GSE31980 | Transcriptome profile in the human synovial MSC-aggregates | 0.66 | 0.64 | 0.60 | 0.63 | 0.66 | -0.71 | 0.62 | 0.70 | 0.71 |
| 211 | GSE45512 | Human Alopecia Areata Skin Profiling | 0.61 | 0.57 | 0.65 | 0.62 | -0.45 | 0.55 | 0.56 | 0.66 | 0.70 |
| 212 | GSE20141 | Expression analysis of laser-dissected SNpc neurons in Parkinson's disease | 0.61 | -0.22 | 0.58 | 0.60 | 0.54 | 0.61 | 0.38 | 0.67 | 0.61 |
| 213 | GSE17032 | Expression data from human fibroblasts | 0.56 | 0.58 | 0.37 | 0.53 | 0.51 | 0.22 | 0.31 | 0.64 | 0.47 |
| 214 | GSE41364 | Expression data for HT29 cells treated with 5-aza-deoxy-cytidine [Affymetrix] | 0.57 | 0.53 | 0.63 | 0.53 | -0.09 | 0.66 | 0.49 | 0.46 | 0.28 |
| 215 | GSE16870 | HeLa cells treated with V-ATPase inhibitors or with desoxyferramine compared to HeLa in DMSO or medium with low LDL | 0.51 | 0.64 | 0.59 | 0.44 | 0.58 | 0.61 | -0.32 | 0.62 | 0.58 |
| 216 | GSE48311 | Time-dependent changes in gene expression after endotoxinic challenge followed by LR12-scrambled or LR12 treatment. | 0.59 | 0.49 | 0.17 | 0.48 | 0.38 | 0.37 | 0.55 | 0.62 | 0.35 |

| | | | | | | | | | | |
|---|---|---|---|---|---|---|---|---|---|---|
| 217 | GSE11142 | Nicotine effect on CEM model T cell line (kuo-affy-human-232861) | 0.61 | 0.38 | -0.07 | 0.59 | 0.56 | 0.32 | 0.63 | 0.60 | 0.62 |
| 218 | GSE49953 | Expression data from two breast cancer cell lines | 0.66 | 0.63 | 0.66 | 0.63 | 0.66 | 0.65 | -0.89 | 0.70 | 0.68 |
| 219 | GSE33630 | Normal thyrocytes vs papillary vs anaplastic thyroid carcinomas | 0.63 | 0.58 | 0.59 | 0.55 | 0.58 | 0.26 | 0.61 | 0.44 | 0.75 |
| 220 | GSE14987 | Expression data from ERBB2 over-expression and EGF stimulation in MCF10A cells | 0.64 | 0.66 | 0.64 | 0.63 | 0.60 | 0.61 | -0.83 | 0.68 | 0.67 |
| 221 | GSE5081 | Expression data from Helicobacter positive and negative human gastritis samples | 0.56 | 0.28 | 0.49 | 0.52 | 0.34 | 0.48 | 0.28 | 0.64 | 0.55 |
| 222 | GSE36765 | Gene expression profiling of CD4+ T cells infiltrating human breast cancer (Discovery Set) | 0.59 | 0.54 | 0.53 | 0.39 | 0.11 | 0.38 | 0.41 | 0.62 | 0.64 |
| 223 | GSE49628 | Large-scale hypomethylated blocks associated with Epstein-Barr virus-induced B-cell immortalization [Expression Array] | 0.66 | 0.50 | 0.55 | 0.61 | 0.47 | 0.61 | -0.38 | 0.70 | 0.65 |
| 224 | GSE12079 | Molecular profiling of CD3- CD4+ T-cells from patients with the lymphocytic variant of hypereosinophilic syndrome | 0.55 | -0.16 | 0.53 | 0.43 | 0.60 | 0.49 | 0.47 | 0.65 | 0.63 |
| 225 | GSE12355 | Detection of Notch1-IC, Notch2-IC and EBNA2 target genes in human B cells | 0.51 | 0.55 | 0.26 | 0.61 | 0.57 | 0.24 | 0.15 | 0.60 | 0.58 |
| 226 | GSE50208 | Molecular-guided therapy predictions reveal drug resistance phenotypes and treatment alternatives in malignant peripheral nerve sheath tumors | 0.63 | 0.64 | 0.60 | 0.62 | 0.61 | -0.38 | 0.17 | 0.69 | 0.60 |
| 227 | GSE4406 | Gene expression profiling of CD4+ T-cells and GM6990 lymphoblastoid cell lines | 0.63 | 0.62 | 0.64 | 0.63 | 0.62 | 0.52 | -0.79 | 0.68 | 0.68 |
| 228 | GSE33325 | Gene expression changes in human cardiomyocytes exposed to VX (O-ethyl S-[2-(diisopropylamino)ethyl] methylphosphonothiolate) | 0.60 | 0.55 | 0.60 | 0.51 | 0.32 | 0.59 | -0.32 | 0.64 | 0.62 |
| 229 | GSE17549 | Loss-of-function mutations in REP-1 affect intracellular vesicle transport in fibroblasts and monocytes of CHM patients | 0.53 | 0.58 | 0.60 | 0.61 | 0.43 | -0.31 | 0.42 | 0.57 | 0.57 |
| 230 | GSE19963 | Expression data from hyperplastic polyps and normal colonic mucosa from patients with familial and sporadic HPPS | 0.59 | 0.51 | 0.44 | 0.60 | 0.38 | 0.61 | -0.29 | 0.64 | 0.46 |
| 231 | GSE18934 | Gene expression in fetal mesenchymal stem cells for identification of epitopes suitable for non-invasive isolation | 0.63 | 0.62 | 0.61 | 0.63 | 0.62 | -0.92 | 0.64 | 0.68 | 0.64 |
| 232 | GSE21483 | Regulation of HB-EGF by miR-212 and acquired cetuximab-resistance in head and neck cancer | 0.58 | 0.61 | 0.62 | 0.63 | 0.62 | 0.64 | -0.91 | 0.44 | 0.64 |
| 233 | GSE16464 | Chondrogenic differentiation potential of OA chondrocytes and their use in autologous chondrocyte transplantation | 0.52 | 0.54 | 0.27 | 0.57 | 0.51 | -0.03 | 0.50 | 0.63 | 0.56 |
| 234 | GSE36767 | Gene changes of CD4+ T cells infiltrating human breast cancer in the absence of tumor environment (Confirmation Set 24h) | 0.64 | 0.63 | 0.57 | 0.58 | 0.42 | -0.65 | 0.60 | 0.67 | 0.69 |
| 235 | GSE18931 | The biological and molecular heterogeneity of breast cancers correlate with their cancer stem cell content | 0.60 | 0.58 | 0.58 | 0.45 | -0.53 | 0.43 | 0.62 | 0.67 | 0.64 |
| 236 | GSE37603 | Identification of WISP1 as an important survival factor in human mesenchymal stem cells | 0.62 | 0.58 | 0.57 | -0.46 | 0.60 | 0.47 | 0.49 | 0.48 | 0.62 |
| 237 | GSE15192 | Differences between CD44+/CD24- and CD44-/CD24+ subpopulation of immortalized human mammary epithelial cells | 0.59 | 0.62 | 0.62 | 0.61 | 0.57 | -0.93 | 0.61 | 0.61 | 0.01 |
| 238 | GSE40241 | Expression data from Versican (VCAN) protein treated ovarian cancer cell line OVCA433 | 0.59 | 0.55 | 0.55 | 0.61 | 0.58 | -0.73 | 0.61 | 0.69 | 0.58 |
| 239 | GSE18912 | Expression profiling of breast cancer cell lines MCF-7 and MCF-7R4 | 0.56 | -0.49 | 0.61 | 0.46 | 0.50 | 0.56 | 0.49 | 0.63 | 0.65 |
| 240 | GSE10289 | Cells silenced for SDHB expression and tumor phenotype | 0.60 | 0.62 | -0.87 | 0.55 | 0.59 | 0.62 | 0.59 | 0.30 | 0.45 |
| 241 | GSE15205 | TGF or TNF Time series in ARPE19 | 0.56 | 0.56 | 0.48 | 0.58 | 0.55 | -0.44 | 0.36 | 0.66 | 0.64 |
| 242 | GSE20948 | The Effect of Hepatitis C Virus Infection on Host Gene Expression | 0.57 | 0.58 | 0.46 | -0.13 | 0.56 | 0.44 | 0.15 | 0.61 | 0.57 |
| 243 | GSE11428 | Expression data from LNCaP and abl cells | 0.52 | 0.33 | 0.27 | 0.37 | 0.48 | 0.44 | 0.22 | 0.61 | 0.48 |
| 244 | GSE35659 | A transcriptional map of the impact of endurance exercise training on skeletal muscle phenotype (resting muscle after endurance training) | 0.39 | 0.50 | 0.50 | 0.12 | 0.34 | 0.20 | 0.55 | 0.48 | 0.58 |
| 245 | GSE31912 | Gene expression profile in MCF7 breast cancer cells after 78 functionallly important molecules were knocked down using siRNA. | 0.51 | 0.54 | 0.45 | 0.49 | 0.37 | 0.31 | -0.05 | 0.54 | 0.33 |

| | | | | | | | | | | |
|---|---|---|---|---|---|---|---|---|---|---|
| 246 | GSE11919 | Vitamin C-induced gene expression profiling in GM5659 human skin fibroblasts | 0.59 | 0.49 | 0.66 | 0.69 | 0.48 | -0.01 | 0.64 | 0.70 | 0.61 |
| 247 | GSE34628 | Gene expression timecourse from Dengue virus infected human endothelial cells | 0.67 | 0.77 | 0.67 | 0.63 | 0.68 | 0.55 | 0.43 | 0.57 | 0.73 |
| 248 | GSE11510 | Taxonomy of placenta cells | 0.55 | 0.58 | 0.54 | 0.48 | 0.35 | 0.46 | -0.41 | 0.64 | 0.32 |
| 249 | GSE17251 | human | 0.57 | 0.53 | 0.57 | 0.07 | 0.35 | 0.48 | -0.03 | 0.59 | 0.58 |
| 250 | GSE21545 | Biobank of Karolinska Endarterectomy (BiKE) | 0.59 | 0.57 | 0.59 | 0.52 | 0.53 | -0.73 | 0.52 | 0.62 | 0.64 |
| 251 | GSE9055 | Time course gene expression of HUVEC after TNF-alpha treatment | 0.50 | 0.53 | 0.48 | 0.15 | 0.50 | 0.04 | 0.31 | 0.46 | 0.49 |
| 252 | GSE35716 | 4HC] | 0.56 | 0.58 | 0.56 | 0.52 | 0.54 | -0.64 | 0.38 | 0.61 | 0.63 |
| 253 | GSE17385 | Gene expression profiling from MM1.S cells with control or beta-catenin knockdown. | 0.61 | -0.62 | 0.09 | 0.59 | 0.60 | 0.60 | 0.61 | 0.66 | 0.66 |
| 254 | GSE30531 | Expression data of A375 melanoma cells after DMSO or MLN4924 treatment from 1 hour to 24 hour | 0.56 | 0.50 | 0.45 | 0.19 | 0.56 | 0.34 | -0.14 | 0.64 | 0.30 |
| 255 | GSE2964 | Motexafin Gadolinium and Zinc Induce Oxidative Stress Responses and Apoptosis in B-Cell Lymphoma Lines | 0.49 | -0.33 | 0.51 | 0.54 | 0.39 | 0.54 | 0.31 | 0.61 | -0.05 |
| 256 | GSE8192 | The DExH-box RNA helicase RHAU is a Nuclear Protein Involved in Transcription and mRNA Decay | 0.38 | 0.52 | 0.52 | 0.37 | 0.44 | 0.57 | -0.39 | 0.61 | 0.48 |
| 257 | GSE4316 | Genome-wide expression profile of human trabecular meshwork cultured cells, non-glaucomatous and POAG tissue | 0.64 | 0.58 | 0.72 | 0.74 | 0.65 | 0.11 | 0.69 | 0.79 | 0.79 |
| 258 | GSE6281 | Gene expression time-course in the human skin during elicitation of allergic contact dermatitis | 0.54 | 0.53 | 0.54 | 0.47 | 0.37 | 0.36 | -0.25 | 0.60 | 0.57 |
| 259 | GSE18995 | Expression data from donor lungs of cardiac death and brain death donors | 0.37 | 0.38 | 0.44 | 0.20 | 0.25 | 0.28 | 0.47 | 0.41 | 0.25 |
| 260 | GSE11430 | AffymetrixDataset | 0.50 | 0.59 | 0.53 | 0.58 | 0.56 | -0.59 | 0.18 | 0.58 | 0.61 |
| 261 | GSE9927 | Chronic CD4+ T cell Activation & Depletion in HIV-1 Infection: Type I Interferon-Mediated Disruption of T Cell Dynamics | 0.45 | 0.37 | -0.11 | 0.43 | 0.52 | 0.43 | 0.25 | 0.54 | 0.47 |
| 262 | GSE10315 | Multipotent mesenchymal stromal cells: identification of pathways common to TGFŒ≤3/BMP2-induced chondrogenesis | 0.46 | 0.51 | 0.45 | 0.34 | 0.45 | -0.29 | 0.41 | 0.54 | 0.61 |
| 263 | GSE14491 | TGFŒ≤/mutant-p53 jointly controlled genes | 0.55 | 0.39 | 0.55 | 0.53 | 0.45 | -0.03 | -0.16 | 0.59 | 0.61 |
| 264 | GSE6519 | Microarray Analysis of Baboon neonates consuming long-chain polyunsaturated fatty acid formula | 0.59 | -0.09 | 0.05 | 0.55 | 0.46 | 0.61 | 0.50 | 0.66 | 0.21 |
| 265 | GSE4107 | Expression profiling in early onset colorectal cancer | 0.19 | 0.48 | 0.42 | 0.39 | 0.41 | -0.13 | 0.48 | 0.06 | 0.44 |
| 266 | GSE11292 | High-time-resolution dynamic analysis of human regulatory T cell (Treg) / CD4+ T-effector cell (Teff) activation | 0.48 | -0.01 | 0.13 | 0.49 | 0.44 | 0.37 | 0.30 | 0.52 | 0.47 |
| 267 | GSE11959 | Anti-IGF-IR antibody h10H5 induces a unique transcriptional profile in SK-N-AS human neuroblastoma xenograft tumor | 0.65 | 0.41 | 0.60 | 0.47 | 0.58 | -0.01 | 0.38 | 0.71 | 0.63 |
| 268 | GSE27838 | Gene expression of expanded and non-expanded natural killer cells from healthy donor and myeloma patients | 0.51 | 0.44 | -0.55 | 0.50 | 0.52 | 0.48 | 0.16 | 0.60 | 0.59 |
| 269 | GSE41828 | TWEAK-treated time course in U2OS cells. | 0.49 | 0.48 | 0.42 | 0.46 | 0.34 | 0.45 | -0.64 | 0.53 | 0.43 |
| 270 | GSE41663 | Re-analysis by microarray using cDNA target of samples from psoriasis patients enrolled in an etanercept trial | 0.54 | 0.30 | 0.50 | 0.45 | -0.08 | 0.51 | -0.23 | 0.57 | 0.55 |
| 271 | GSE44029 | Expression data from SW480 cells with Gankyrin knockdown | 0.52 | 0.04 | 0.53 | 0.49 | -0.66 | 0.52 | 0.54 | 0.54 | 0.55 |
| 272 | GSE23103 | HeLa SCY1-like 1 esiRNA knockdown | 0.60 | 0.60 | 0.59 | 0.59 | 0.57 | 0.55 | -0.90 | 0.67 | 0.64 |
| 273 | GSE39059 | Changes in microRNA and mRNA expression with differentiation of human bronchial epithelial cells [mRNA] | 0.14 | 0.50 | 0.55 | 0.50 | 0.52 | 0.53 | -0.77 | -0.26 | 0.59 |
| 274 | GSE28005 | Charaterization of the initial molecular events of adipose tissue development and growth during overfeeding in humans | 0.54 | 0.53 | 0.53 | 0.42 | 0.23 | 0.10 | 0.25 | 0.60 | 0.55 |
| 275 | GSE48350 | Alzheimer's Disease Dataset | 0.47 | -0.18 | 0.45 | 0.35 | 0.34 | 0.41 | 0.07 | 0.51 | 0.41 |
| 276 | GSE30188 | Rho transcription inhibitor CCG-1423 effect on PC-3 cells | 0.51 | 0.49 | -0.66 | 0.51 | 0.48 | 0.51 | 0.06 | 0.59 | 0.55 |
| 277 | GSE37364 | Expression data from human colonic biopsy samples (adenoma-carcinoma) | 0.23 | 0.24 | 0.10 | 0.46 | 0.25 | 0.43 | 0.24 | 0.37 | 0.19 |

| | | | | | | | | | | |
|---|---|---|---|---|---|---|---|---|---|---|
| 278 | GSE18271 | Analysis of TALE homeobox genes in neuroblastic tumors: ganglioneuroblastoma and ganglioneuroma | 0.48 | -0.53 | 0.50 | 0.41 | 0.39 | 0.41 | 0.19 | 0.56 | 0.38 |
| 279 | GSE47855 | Gene expression analysis for CD56- T, NK, CD56+ T cells, and iNKT cells | 0.35 | 0.35 | -0.01 | 0.32 | 0.34 | 0.39 | 0.10 | 0.54 | 0.56 |
| 280 | GSE10718 | Time course of NHBE cells exposed to whole cigarette smoke (full flavor) | 0.43 | 0.48 | -0.15 | 0.21 | 0.39 | 0.42 | 0.01 | 0.48 | 0.27 |
| 281 | GSE7637 | Expression data from human mesenchymal stem cells (#4F1560) | 0.43 | 0.46 | 0.49 | 0.28 | 0.47 | -0.50 | 0.18 | 0.57 | 0.47 |
| 282 | GSE20297 | The effects of terbutaline or GW9508 on TNF-alpha and IFN gamma (TNF-alpha + IFN gamma) stimulation by HaCaT | 0.49 | 0.47 | 0.45 | 0.47 | 0.11 | 0.48 | -0.71 | 0.57 | 0.56 |
| 283 | GSE10879 | Expression data of hormone-responsive MCF-7 cells versus estrogen-deprived MCF-7:5C and MCF-7:2A breast cancer cells | 0.34 | 0.52 | -0.04 | 0.49 | 0.40 | -0.45 | 0.50 | 0.40 | 0.50 |
| 284 | GSE47685 | Gene silencing of BSK65-MONO1 (RNF185) and of its natural antisense RNA (RNF185-AS) using siRNAs | 0.66 | 0.57 | 0.67 | 0.67 | -0.21 | 0.58 | 0.46 | 0.65 | 0.43 |
| 285 | GSE8671 | Transcriptome profile of human colorectal adenomas. | 0.44 | 0.12 | 0.22 | 0.44 | 0.37 | 0.47 | 0.04 | 0.46 | 0.33 |
| 286 | GSE13070 | Human Insulin Resistance and Thiazolidinedione-Mediated Insulin Sensitization | 0.46 | 0.44 | 0.45 | 0.00 | 0.44 | -0.46 | 0.41 | 0.54 | 0.53 |
| 287 | GSE33950 | SHARP1 suppresses breast cancer metastasis by promoting degradation of hypoxia-inducible factors | 0.40 | 0.50 | 0.49 | 0.37 | 0.25 | -0.80 | 0.49 | 0.34 | 0.22 |
| 288 | GSE17612 | Comparison of post-mortem tissue from brain BA10 region between schizophrenic and control patients. | 0.37 | -0.30 | 0.41 | 0.31 | 0.20 | 0.36 | 0.28 | 0.44 | 0.36 |
| 289 | GSE49910 | An Expression Atlas of Human Primary Cells: Inference of Gene Function from Coexpression Networks | 0.50 | 0.44 | 0.43 | 0.46 | 0.44 | -0.16 | -0.44 | 0.57 | 0.53 |
| 290 | GSE41296 | Characterization of Formaldehyde's Genotoxic Mode of Action by Gene Expression Analysis in TK6 Cells | 0.45 | -0.19 | 0.19 | 0.45 | 0.43 | 0.38 | -0.11 | 0.53 | 0.51 |
| 291 | GSE42046 | TWEAK-treated time course in ACHN cells | 0.49 | 0.45 | 0.32 | 0.40 | 0.42 | 0.27 | -0.78 | 0.57 | 0.45 |
| 292 | GSE10311 | Systematic Assessment of Human Osteoblast Transcriptome in Resting and Induced Primary Cells | 0.62 | 0.61 | 0.60 | 0.08 | 0.64 | 0.08 | 0.61 | 0.69 | 0.61 |
| 293 | GSE15918 | Torcetrapib induces aldosterone and cortisol production in an intracellular calcium-dependent mechanism | 0.45 | 0.42 | 0.47 | -0.11 | -0.47 | 0.46 | 0.32 | 0.53 | 0.14 |
| 294 | GSE11238 | Vaccinia E3L mutant virus infected HeLa cell lines (langl-affy-human-215499) | 0.42 | 0.35 | 0.06 | 0.46 | 0.45 | -0.46 | 0.23 | 0.53 | 0.15 |
| 295 | GSE12548 | EMT Time series in ARPE19 | 0.38 | 0.47 | 0.39 | 0.39 | 0.45 | -0.60 | 0.15 | 0.56 | 0.43 |
| 296 | GSE20318 | YWHAZ is an Invasion and Metastasis promoting genes of Lung cancer | 0.76 | 0.72 | 0.51 | 0.72 | 0.68 | -0.09 | 0.74 | 0.78 | 0.45 |
| 297 | GSE15013 | Expression of HOXB genes is significantly different in acute myeloid leukemia with a partial tandem duplication of MLL vs. a MLL translocation: a cross-laboratory study | 0.39 | -0.11 | 0.39 | 0.40 | 0.31 | 0.19 | -0.18 | 0.47 | 0.51 |
| 298 | GSE3202 | MK886 treatment of H720 non-small cell lung cancer cell line | 0.42 | -0.04 | 0.24 | 0.42 | 0.46 | -0.21 | 0.29 | 0.49 | 0.23 |
| 299 | GSE32473 | Gene expression is differently affected by pimecrolimus and betamethasone in lesional skin of atopic dermatitis. | 0.36 | 0.37 | 0.40 | 0.35 | -0.15 | 0.37 | -0.34 | 0.50 | 0.36 |
| 300 | GSE47751 | Early tissue responses to etanercept in psoriasis lesions | 0.43 | 0.22 | 0.38 | 0.36 | -0.04 | 0.38 | -0.38 | 0.47 | 0.39 |
| 301 | GSE33050 | GlcNAcylation of histone H2B facilitates its monoubiquitination [Affymetrix data] | 0.43 | 0.39 | 0.39 | 0.49 | 0.24 | 0.45 | -0.80 | 0.51 | -0.05 |
| 302 | GSE13564 | Gene expression in the human prefrontal cortex during postnatal development | 0.43 | -0.25 | 0.42 | 0.31 | 0.36 | -0.34 | 0.37 | 0.51 | 0.45 |
| 303 | GSE5563 | Gene expression profile of VIN lesions in comparison to controls | 0.46 | 0.22 | 0.36 | 0.33 | 0.36 | 0.39 | -0.54 | 0.50 | 0.49 |
| 304 | GSE32876 | Inferring transcriptional and microRNA-mediated regulatory programs in glioblastoma | 0.35 | 0.39 | -0.16 | 0.09 | 0.41 | 0.33 | -0.25 | 0.42 | 0.40 |
| 305 | GSE8056 | Gene Expression Profiles in Thermally Injured Human Skin: A Temporal Microarray Analysis | 0.45 | 0.41 | 0.49 | 0.47 | 0.52 | 0.37 | -0.52 | 0.59 | 0.41 |
| 306 | GSE27390 | Human bone marrow-derived mononuclear cells (BMMC): rheumatoid arthritis vs. osteoarthritis | 0.37 | 0.38 | 0.39 | 0.35 | 0.30 | -0.58 | -0.14 | 0.46 | 0.33 |
| 307 | GSE15132 | Riboflavin depletion impairs cell proliferation in intestinal cells: Identification of mechanisms and consequences | 0.38 | 0.45 | 0.22 | 0.37 | 0.35 | -0.20 | -0.27 | 0.50 | -0.02 |
| 308 | GSE44765 | Global profiling of human hair follicle scalp dermal papilla cells using Affymetrix microarrays | 0.75 | 0.69 | 0.69 | 0.42 | 0.76 | 0.70 | 0.41 | 0.80 | 0.82 |

| | | | | | | | | | | |
|---|---|---|---|---|---|---|---|---|---|---|
| 309 | GSE51524 | LNCaP prostate cancer cell lines overexpressing wild-type or GARRPR-mutant Bag-1L | 0.38 | -0.56 | 0.37 | 0.40 | 0.39 | -0.36 | 0.35 | 0.47 | 0.47 |
| 310 | GSE32526 | Expression data from breast cancer tumor-initiating cells | 0.40 | 0.40 | 0.28 | 0.40 | 0.40 | -0.28 | -0.66 | 0.43 | 0.17 |
| 311 | GSE8687 | Inhibition of activation of Sez-4 cell line with IL-2 by Jak kinase inhibitors. | 0.32 | 0.29 | 0.28 | 0.36 | -0.17 | 0.30 | -0.47 | 0.33 | 0.13 |
| 312 | GSE7216 | Cytokine treated normal human epidermal keratinocytes | 0.28 | 0.07 | 0.28 | 0.18 | 0.01 | 0.37 | -0.32 | 0.43 | 0.31 |
| 313 | GSE3151 | Oncogene Signature Dataset | 0.16 | 0.22 | 0.25 | 0.06 | -0.02 | 0.07 | 0.12 | 0.29 | 0.28 |
| 314 | GSE12293 | Evolution of neuronal and endothelial transcriptomes in primates | 0.37 | -0.42 | 0.36 | 0.30 | -0.38 | 0.33 | 0.29 | 0.44 | 0.41 |
| 315 | GSE49353 | Evaluating cross-hybridization of murine cDNA to the Affymetrix Human Genome U133 Plus 2.0 chipset | -0.33 | 0.25 | 0.25 | 0.34 | 0.32 | -0.45 | 0.33 | -0.09 | -0.19 |
| 316 | GSE15520 | The Role of Cholesterol Pathways in Norovirus Replication | 0.28 | 0.53 | 0.42 | 0.50 | 0.54 | 0.44 | -0.77 | 0.28 | -0.13 |
| 317 | GSE22779 | Gene expression data of non-leukemic individuals before and during in-vivo glucocorticoid treatment | 0.25 | -0.02 | 0.01 | 0.23 | 0.24 | -0.16 | 0.02 | 0.36 | 0.38 |
| 318 | GSE31681 | Human cumulus cells | 0.30 | 0.28 | 0.23 | 0.06 | 0.00 | -0.38 | 0.23 | 0.40 | 0.27 |
| 319 | GSE34599 | In-transit extremity melanoma III | 0.54 | 0.50 | 0.52 | 0.26 | 0.18 | 0.21 | 0.35 | 0.60 | 0.53 |
| 320 | GSE13987 | Profile of rolipram treated B-CLL, normal B, and normal T cells | 0.28 | 0.25 | 0.30 | -0.37 | -0.54 | 0.28 | 0.26 | 0.39 | 0.37 |
| 321 | GSE24337 | The Human Airway Epithelial Basal Cell Transcriptome | -0.14 | 0.19 | -0.07 | 0.21 | 0.26 | 0.22 | -0.24 | 0.03 | 0.25 |
| 322 | GSE10739 | LPS and PMA response in parental MM6 cells | 0.34 | 0.33 | -0.64 | 0.33 | 0.33 | 0.33 | -0.55 | 0.44 | -0.25 |
| 323 | GSE11864 | Effect of interferon-gamma on macrophage differentiation and response to Toll-like receptor ligands | 0.32 | 0.31 | 0.20 | 0.20 | 0.28 | -0.47 | -0.53 | 0.42 | 0.37 |
| 324 | GSE29368 | CD140a+ human oligodendrocyte progenitor cells | 0.29 | -0.40 | 0.23 | -0.48 | 0.25 | 0.08 | 0.25 | 0.33 | 0.36 |
| 325 | GSE46873 | Dual targeting of MYC and CYCLON by BET bromodomain inhibition optimizes Rituximab response in lymphoma. | 0.45 | 0.38 | -0.58 | 0.26 | 0.41 | 0.43 | 0.31 | 0.37 | 0.21 |
| 326 | GSE15824 | Gene expression profiling of human gliomas and human glioblastoma cell lines | 0.26 | 0.24 | -0.03 | 0.28 | 0.30 | 0.19 | -0.44 | 0.39 | 0.23 |
| 327 | GSE8139 | Expression data from MCF7/HER2-18 xenografts | 0.03 | 0.10 | -0.03 | -0.04 | 0.06 | -0.04 | 0.13 | 0.13 | 0.21 |
| 328 | GSE33585 | Expression data from monocytic cell lines (THP) | 0.25 | 0.27 | -0.39 | 0.25 | 0.05 | 0.26 | -0.58 | 0.38 | 0.35 |
| 329 | GSE19278 | 2] | 0.41 | 0.29 | -0.06 | 0.36 | 0.34 | 0.31 | -0.51 | 0.48 | 0.48 |
| 330 | GSE9517 | Cysteine deprivation in liver cell line | 0.33 | 0.08 | 0.32 | 0.03 | 0.02 | 0.37 | -0.10 | 0.31 | 0.33 |
| 331 | GSE52158 | Dynamic developmental signaling logic underlying lineage bifurcations during human endoderm induction and patterning from pluripotent stem cells [Expression data set] | 0.21 | 0.14 | 0.22 | 0.08 | -0.38 | 0.15 | -0.33 | 0.31 | 0.15 |
| 332 | GSE24468 | Elucidation of the Mechanisms by which the Progesterone Receptor Inhibits Inflammatory Responses in Cellular Models of Breast Cancer | 0.12 | -0.20 | -0.09 | -0.03 | -0.02 | -0.04 | 0.01 | 0.27 | 0.23 |
| 333 | GSE10070 | Gene Expression in MCF10A cells through Differentiation on Transwells | 0.32 | 0.29 | -0.41 | 0.23 | 0.26 | 0.31 | -0.36 | 0.36 | 0.36 |
| 334 | GSE18235 | Effect of 10 Cigarette Smoke Condensates on Primary Human Airway Epithelial Cells | 0.44 | 0.40 | -0.32 | 0.35 | 0.38 | 0.45 | 0.02 | 0.47 | 0.30 |
| 335 | GSE23610 | Gene expression profiles of MCF-7 cells treated with Si-Wu-Tang, estradiol and ferulic acid | 0.08 | -0.32 | -0.32 | -0.02 | 0.04 | -0.18 | 0.06 | 0.17 | 0.17 |
| 336 | GSE36701 | Gene expression analysis of rectal mucosa in chronic irritable bowel syndrome (IBS) compared to healthy volunteers (HV) | 0.39 | 0.52 | 0.54 | 0.57 | 0.18 | 0.60 | 0.53 | 0.65 | 0.62 |
| 337 | GSE17400 | Dynamic Innate Immune Responses of Human Bronchial Epithelial Cells against SARS-CoV and DOHV infection | 0.35 | 0.64 | 0.29 | 0.50 | 0.59 | 0.66 | 0.59 | 0.15 | 0.39 |
| 338 | GSE40266 | Expression data from TGF-beta-treated human ovarian fibroblasts | 0.04 | -0.33 | -0.19 | 0.03 | 0.08 | -0.37 | 0.09 | 0.15 | 0.02 |
| 339 | GSE11941 | Topoisomerase II inhibition involves characteristic chromosomal expression patterns: Trovafloxacin study | 0.65 | 0.73 | 0.57 | 0.52 | 0.74 | 0.70 | 0.47 | 0.79 | 0.81 |
| 340 | GSE53603 | Expression data from SKOV3 cells treated with SAHA or vehicle control | 0.61 | 0.51 | 0.63 | 0.58 | 0.43 | 0.29 | -0.09 | 0.59 | 0.49 |
| 341 | GSE10281 | Letrozole (Femara) early response to treatment | 0.50 | 0.48 | 0.44 | 0.07 | 0.37 | 0.50 | 0.42 | 0.42 | 0.52 |

| | | | | | | | | | | | |
|---|---|---|---|---|---|---|---|---|---|---|---|
| 342 | GSE8784 | Plasmodium Circumsporozoite Protein Promotes the Development of the Liver Stages of the Parasite | 0.28 | -0.29 | 0.19 | 0.30 | 0.18 | 0.29 | -0.43 | 0.42 | 0.36 |
| 343 | GSE32967 | Modeling lethal prostate cancer variant with small cell carcinoma features [expression profile] | 0.65 | 0.32 | 0.28 | 0.61 | 0.47 | 0.54 | 0.63 | 0.71 | 0.72 |
| 344 | GSE25619 | Gene expression profiles of granulin- and control-treated normal human mammary fibroblasts | 0.72 | 0.66 | 0.66 | 0.26 | 0.59 | 0.53 | 0.41 | 0.72 | 0.74 |
| 345 | GSE16480 | Inactivation of CDK2 is synthetic lethal to MYCN-overexpressing cancer cells | 0.46 | 0.56 | 0.44 | 0.55 | 0.53 | 0.54 | -0.05 | 0.63 | 0.41 |
| 346 | GSE9984 | Profiling Gene Expression in Human Placentae of Different Gestational Ages: an OPRU Network and UW SCOR Study | 0.64 | 0.64 | 0.36 | 0.64 | 0.65 | 0.16 | 0.44 | 0.63 | 0.69 |
| 347 | GSE33495 | Disrupted transcripitonal network in ŒîNp63 AEC tissue model [gene expression] | 0.67 | 0.60 | 0.28 | 0.61 | 0.07 | 0.64 | 0.24 | 0.67 | 0.66 |
| 348 | GSE12963 | Gene expression in human CD4+ T-lymphocytes infected with VSVG-pseudotyped HIV-1 viruses lacking Env, Vpr, and Nef | 0.62 | 0.56 | -0.14 | 0.53 | 0.45 | 0.55 | 0.62 | 0.63 | 0.49 |
| 349 | GSE9250 | Genomic profiling in CLL and subtypes of del13q14 | 0.63 | 0.21 | 0.46 | 0.50 | 0.51 | 0.35 | 0.56 | 0.63 | 0.59 |
| 350 | GSE31215 | Gene expression analysis of human pediatric mesenchymal stem cells (hpMSCs) upon expression of EWS-FLI-1 | 0.45 | 0.52 | 0.56 | 0.49 | 0.56 | 0.26 | 0.44 | 0.51 | 0.43 |
| 351 | GSE4737 | HCaRG vs NEO | 0.69 | 0.70 | 0.66 | 0.19 | 0.54 | 0.51 | 0.63 | 0.71 | 0.63 |
| 352 | GSE30439 | Exposure of cystic fibrosis bronchial epithelial cells (CFBE 41 o-) to Pseudomonas aeruginosa (PA01) biofilms | 0.63 | 0.54 | 0.27 | 0.60 | 0.55 | 0.57 | 0.09 | 0.64 | 0.60 |
| 353 | GSE40986 | Gene expression profiles induced by overexpression of PDEF in MCF10A mammary epithelial cell line | 0.56 | -0.39 | 0.15 | 0.62 | 0.58 | 0.62 | 0.59 | 0.52 | 0.48 |
| 354 | GSE38718 | Sex and aging effect on skeletal muscle transcriptome in humans | 0.38 | 0.39 | 0.37 | -0.08 | 0.40 | -0.16 | 0.30 | 0.40 | 0.40 |
| 355 | GSE33112 | Gene expression in colon cancer stem cells (CSC) cultures identified by Wnt signaling levels | 0.62 | 0.59 | 0.67 | 0.62 | 0.65 | 0.27 | 0.36 | 0.71 | 0.48 |
| 356 | GSE8640 | TFAP2C regulates multiple pathways of estrogen signaling | 0.45 | 0.56 | 0.37 | 0.35 | 0.35 | 0.41 | -0.17 | 0.60 | 0.57 |
| 357 | GSE22148 | Induced Sputum Genes Associated With Spirometroc and Radiological Disease Severity in COPD Ex-smokers | 0.43 | 0.38 | 0.40 | 0.43 | 0.36 | -0.22 | 0.06 | 0.50 | 0.50 |
| 358 | GSE57552 | ZFX silencing introduced differential gene expression in leukemia cells | 0.47 | 0.48 | 0.45 | 0.30 | -0.01 | 0.48 | -0.80 | 0.53 | 0.54 |
| 359 | GSE24530 | Identification and Characterization of Subpopulations within Human Embryonic Stem Cell Lines | 0.65 | 0.68 | 0.70 | 0.52 | 0.37 | 0.30 | 0.59 | 0.75 | 0.61 |
| 360 | GSE2817 | Wavelet modelling of microarray data provides chromosomal pattern of expression which predicts survival in gliomas | 0.43 | 0.27 | 0.27 | 0.38 | 0.16 | 0.10 | 0.43 | 0.57 | 0.45 |
| 361 | GSE29330 | Identification of GNG7 as An Epigenetically Silenced Gene in Head and Neck Cancer by Gene Expression Profiling | 0.59 | 0.57 | 0.60 | 0.26 | 0.25 | 0.53 | 0.23 | 0.54 | 0.56 |
| 362 | GSE33424 | Expression data from human cord blood CD161++/CD161+/CD161- CD8+ T cell subsets | 0.45 | 0.13 | 0.45 | 0.09 | -0.51 | 0.42 | 0.40 | 0.54 | 0.47 |
| 363 | GSE41035 | FGFR3-shRNA induced transcriptional changes in RT112 bladder cancer cells | 0.51 | -0.11 | 0.25 | 0.49 | 0.48 | 0.54 | 0.15 | 0.60 | 0.59 |
| 364 | GSE31782 | Knock-down and Over-expression of JMJD6 in MCF-7 and/or MDA-MB231 | 0.22 | 0.52 | 0.52 | 0.47 | 0.47 | 0.44 | -0.38 | 0.29 | -0.08 |
| 365 | GSE13274 | Ad-HER-wt and Ad-HER2-ki infected HMECs | 0.51 | 0.41 | 0.50 | 0.12 | 0.45 | 0.40 | -0.07 | 0.54 | 0.57 |
| 366 | GSE36287 | Expression data from primary human keratinocytes exposed to cytokines in vitro (IL-4, IL-13, IL-17A, IFN-alpha, IFN-gamma, TNF) | 0.47 | 0.60 | 0.52 | 0.53 | 0.22 | 0.60 | 0.32 | 0.64 | 0.21 |
| 367 | GSE16237 | Expression data of human neuroblastoma tissue samples | 0.51 | 0.11 | 0.47 | 0.48 | 0.52 | 0.19 | 0.44 | 0.63 | 0.47 |
| 368 | GSE21979 | Transcriptional and post-transcriptional regulation of VEGF by the unfolded protein response | 0.63 | 0.60 | 0.43 | 0.56 | 0.56 | 0.60 | -0.23 | 0.61 | 0.53 |
| 369 | GSE39902 | Role of TAZ as mediator of Wnt signaling (MII) | 0.55 | 0.52 | 0.59 | 0.24 | 0.54 | 0.49 | -0.13 | 0.48 | 0.55 |
| 370 | GSE8685 | Activation of Sez-4 cell line with IL-2, IL-15 or IL-21. | 0.20 | 0.31 | 0.26 | 0.28 | -0.21 | 0.34 | -0.45 | 0.43 | 0.40 |
| 371 | GSE6140 | Cross platform microarray analysis for robust identification of differentially expressed genes | 0.01 | 0.73 | 0.74 | 0.73 | 0.70 | 0.66 | 0.70 | -0.41 | -0.34 |
| 372 | GSE14668 | B-Cell Gene Signature with Massive Intrahepatic Production of Antibodies to | 0.55 | 0.41 | 0.43 | 0.60 | 0.43 | 0.57 | 0.39 | 0.63 | 0.65 |

| | | | | | | | | | | | |
|---|---|---|---|---|---|---|---|---|---|---|---|
| | | Hepatitis B Core Antigen in HBV-Associated Acute Liver Failure | | | | | | | | | |
| 373 | GSE11367 | Effect of IL-17 on human vascular smooth muscle cells | 0.62 | 0.56 | 0.54 | 0.12 | 0.42 | 0.58 | 0.32 | 0.66 | 0.72 |
| 374 | GSE21668 | Expression data from undifferentiated human embryonic stem cells (hESC) and Day 3.5 mesodermal progenitor (CD326neg CD56+) population | 0.31 | 0.72 | 0.61 | 0.76 | 0.70 | 0.55 | 0.68 | 0.79 | 0.75 |
| 375 | GSE24422 | Effect of insulin on the stromal and adipocyte cells within hMSC derived adipocytes | 0.60 | 0.43 | 0.42 | 0.18 | 0.31 | 0.56 | 0.04 | 0.63 | 0.56 |
| 376 | GSE41802 | Isocitrata Dehydrogenase (IDH) Mutations Promote a Reversible ZEB1/mir-200-Dependent Epithelial Mesenchymal Transition (EMT) | 0.42 | 0.60 | 0.47 | 0.50 | 0.35 | 0.44 | 0.45 | 0.63 | 0.42 |
| 377 | GSE5486 | Using GINI2 to identify novel mutations in candidate tumor suppressor genes in colon cancer cells | 0.60 | 0.50 | 0.38 | 0.07 | 0.44 | 0.41 | 0.49 | 0.42 | 0.23 |
| 378 | GSE19735 | Comparison of human embroynic stem cell derived vascular cells to mature human vascular and hematopoietic cells | 0.55 | 0.48 | 0.37 | 0.54 | -0.05 | -0.25 | 0.55 | 0.59 | 0.53 |
| 379 | GSE7874 | Effects of EPO and EST on erythroid maturation | -0.26 | 0.16 | 0.11 | 0.17 | 0.20 | -0.49 | 0.17 | 0.15 | 0.25 |
| 380 | GSE10046 | Breast cancer-associated fibroblasts confer AKT1-mediated epigenetic silencing of Cystatin M in epithelial cells. | 0.77 | 0.82 | 0.65 | 0.59 | 0.57 | 0.76 | 0.60 | 0.83 | 0.75 |
| 381 | GSE30127 | Establishment of human trophoblast progenitor cell lines from the chorion | 0.62 | 0.06 | 0.31 | 0.59 | 0.64 | 0.60 | 0.31 | 0.56 | 0.61 |
| 382 | GSE40873 | a prospective study | 0.54 | 0.08 | 0.55 | 0.32 | 0.48 | 0.48 | 0.15 | 0.59 | 0.60 |
| 383 | GSE11618 | Stable XIAP knockdown in HCT116 colon cancer cells | 0.57 | 0.46 | 0.27 | 0.58 | 0.59 | 0.50 | 0.11 | 0.65 | 0.66 |
| 384 | GSE30660 | The Effect of Repeated Whole Cigarette Smoke Challenge on Human Air-Liquid Interface Lung Epithelial Cultures | -0.40 | 0.26 | 0.22 | 0.21 | 0.32 | 0.33 | -0.53 | -0.24 | -0.04 |
| 385 | GSE9169 | Gene expression during neuronal differentiation in two subtypes of SH-SY5Y | 0.51 | 0.26 | 0.41 | 0.52 | 0.43 | 0.39 | 0.25 | 0.64 | 0.54 |
| 386 | GSE44807 | Gene expression data from primary human bronchial epithelial cells expressing EGFP or DN-GRHL2 | 0.61 | 0.50 | 0.60 | -0.30 | 0.58 | 0.04 | 0.61 | 0.18 | 0.65 |
| 387 | GSE7586 | Genome wide analysis of placental malaria | 0.38 | 0.52 | 0.47 | 0.06 | 0.45 | -0.20 | 0.53 | 0.46 | 0.56 |
| 388 | GSE53552 | Gene expression profiling in psoriatic lesional and non-lesional skin [brodalumab treatment] | 0.50 | 0.22 | 0.48 | 0.34 | 0.14 | 0.44 | -0.09 | 0.55 | 0.51 |
| 389 | GSE40281 | Signaling pathways of HPAIV | 0.55 | 0.61 | 0.61 | 0.55 | 0.31 | 0.61 | 0.39 | 0.64 | 0.21 |
| 390 | GSE15773 | Expression data from human adipose tissue | 0.52 | 0.42 | 0.46 | 0.44 | 0.11 | -0.24 | 0.39 | 0.58 | 0.44 |
| 391 | GSE39454 | Genomic signatures characterize leukocyte infiltration in myositis muscles | 0.48 | 0.42 | 0.47 | -0.01 | 0.27 | -0.28 | 0.37 | 0.44 | 0.55 |
| 392 | GSE41386 | Role of REST in the pathogenesis of uterine fibroids | 0.59 | 0.69 | 0.40 | 0.67 | 0.45 | 0.63 | 0.45 | 0.63 | 0.63 |
| 393 | GSE55529 | EcadEGFP expression in MDA-MB-134 and IPH-926 | 0.68 | 0.54 | 0.34 | 0.61 | 0.46 | 0.61 | 0.59 | 0.58 | 0.77 |
| 394 | GSE40730 | Genome-wide analysis of RNAs translationally regulated upon BRCA1 depletion in human mammary epithelial cells | 0.66 | 0.67 | 0.62 | 0.12 | 0.31 | 0.63 | 0.53 | 0.66 | -0.27 |
| 395 | GSE27128 | Expression levels in strained vs. non-strained Calu-3 lung epithelial cells | 0.57 | 0.61 | 0.60 | 0.49 | 0.53 | 0.54 | -0.14 | 0.52 | 0.13 |
| 396 | GSE32100 | Glioma cells oxygen response | 0.06 | -0.22 | -0.07 | -0.13 | 0.05 | -0.21 | -0.09 | 0.10 | 0.10 |
| 397 | GSE4600 | Identifying targets of MeCP2 during neuronal maturational differentiation | 0.48 | -0.38 | 0.22 | 0.41 | 0.49 | 0.47 | 0.53 | 0.54 | 0.49 |
| 398 | GSE10595 | Interaction of bone marrow stroma and monocytes: bone marrow stromal cell lines cultured with monocytes | 0.69 | 0.63 | 0.64 | 0.67 | 0.61 | -0.47 | 0.51 | 0.73 | 0.01 |
| 399 | GSE25087 | Human Fetal and Adult Peripheral Na√Øve CD4+ T cells and CD4+CD25+ Treg cells | 0.57 | 0.47 | 0.31 | 0.44 | 0.51 | -0.45 | 0.45 | 0.63 | 0.61 |
| 400 | GSE39843 | Expression data of cystic fibrosis and non-cystic fibrosis airway cell lines under oxidative stress | 0.64 | 0.58 | 0.45 | 0.29 | 0.46 | 0.20 | 0.45 | 0.66 | 0.68 |
| 401 | GSE10575 | Migratory chondrogenic progenitor cells from repair tissue during the later stages of human osteoarthritis | 0.54 | 0.52 | 0.37 | 0.24 | 0.50 | 0.20 | -0.19 | 0.62 | 0.46 |
| 402 | GSE5675 | Pilocytic astrocytoma | 0.42 | 0.35 | 0.45 | 0.38 | 0.40 | 0.20 | 0.45 | 0.61 | 0.56 |
| 403 | GSE23640 | Gene-expression profile of breast cancer cell lines and sorted breast cancer epithelial cells | 0.50 | 0.55 | 0.29 | 0.30 | 0.52 | 0.54 | 0.20 | 0.48 | 0.61 |
| 404 | GSE4975 | Expression data from p63 siRNA in squamous cell lines | 0.59 | 0.58 | 0.62 | 0.36 | -0.45 | 0.63 | 0.38 | 0.68 | 0.28 |

| | | | | | | | | | | |
|---|---|---|---|---|---|---|---|---|---|---|
| 405 | GSE40968 | The effect of ACSL4 expression on overall gene expression in breast cancer cell lines | 0.68 | 0.61 | 0.62 | 0.55 | 0.31 | 0.48 | 0.50 | 0.70 | 0.66 |
| 406 | GSE10580 | Genes regulated by PRDM5 in U2OS cells. | 0.53 | 0.50 | -0.24 | 0.57 | 0.42 | 0.30 | 0.57 | 0.56 | 0.26 |
| 407 | GSE34828 | Expression data from fibroblast growth factor receptor 4 (FGFR4) knock down ovarian cancer cell lines | 0.66 | 0.70 | 0.67 | 0.24 | 0.35 | 0.34 | 0.65 | 0.61 | 0.32 |
| 408 | GSE20086 | Heterogeneity of gene expression in stromal fibroblasts of human breast carcinomas and normal breast | 0.63 | 0.30 | 0.52 | 0.36 | 0.39 | 0.51 | 0.53 | 0.67 | 0.64 |
| 409 | GSE11729 | H1299 EGF and Iressa stimulation | 0.49 | 0.55 | 0.47 | 0.53 | 0.39 | 0.19 | 0.05 | 0.49 | 0.11 |
| 410 | GSE50175 | Expression data from human Th1 and Th1Th17 cells | 0.63 | 0.06 | 0.40 | 0.64 | 0.55 | 0.61 | 0.33 | 0.66 | 0.69 |
| 411 | GSE43177 | MicroRNA regulate immunological pathways in T-cells in immune thrombocytopenia (ITP) [mRNA] | 0.68 | 0.16 | 0.47 | 0.66 | 0.32 | 0.67 | 0.66 | 0.71 | 0.71 |
| 412 | GSE41485 | Expression data of A939572 SCD1 inhibitor treated ccRCC cells | 0.54 | 0.48 | 0.33 | 0.42 | 0.69 | 0.52 | 0.54 | 0.74 | 0.37 |
| 413 | GSE35006 | Profiling of p53-responsive genes in human breast cancer cells harboring endogenous ts-p53 E285K | 0.73 | 0.53 | 0.20 | 0.65 | 0.66 | 0.58 | 0.63 | 0.70 | 0.70 |
| 414 | GSE14474 | The Effects of Static Magnetic Fields on Human Embryonic Cells | 0.33 | 0.54 | 0.60 | 0.19 | 0.62 | 0.28 | 0.51 | 0.62 | 0.54 |
| 415 | GSE8507 | Neutrophil and PBMC gene expression data from Job's Syndrome | 0.38 | 0.38 | 0.39 | 0.27 | 0.33 | 0.00 | -0.15 | 0.52 | 0.25 |
| 416 | GSE9101 | Expression data in native lipoprotein-stimulated human THP-1 macrophages | 0.29 | 0.36 | 0.28 | -0.31 | -0.43 | 0.36 | 0.32 | 0.44 | 0.38 |
| 417 | GSE40215 | shRNA knockdown of the transcription factor NF-YA (NFYA) | 0.52 | 0.51 | 0.54 | 0.51 | 0.35 | 0.54 | -0.82 | 0.59 | 0.56 |
